# Artificial Intelligence-driven Whole-brain Cell Mapping with Highly Multiplexed In Situ Hybridization

**DOI:** 10.1101/2025.08.18.670857

**Authors:** Tatsuya C. Murakami, Meng Xia, Yurie Maeda, Yuejia Yin, Paolo Emilio Barbano, Ziyi Lin, Tomoyuki Mano, Kazuki Tainaka, Sam Reiter, Nathaniel Heintz

**Affiliations:** Laboratory of Molecular Biology, The Rockefeller University, 1230 York Avenue, New York, NY 10065, USA; Data Science Platform, The Rockefeller University, 1230 York Avenue, New York, NY 10065, USA; Independent researcher; Computational Neuroethology Unit, Okinawa Institute of Science and Technology (OIST) Graduate University, Okinawa 904-0495, Japan; System Pathology for Neurological Disorders, Brain Research Institute, Niigata University, Niigata 951-8122, Japan

## Abstract

Recent advances in three-dimensional single-cell-resolution imaging have begun to link organ-wide and cellular level research in development and disease research. Harnessing the power of whole-mount cell staining and tissue-clearing, it became possible to quantify the cell populations throughout an intact organ. While powerful, whole-organ imaging remains limited by the inability to stain a broad range of molecular markers simultaneously and by the lack of an analytical scheme to precisely quantify the cell population. Here, we present a highly multiplexed whole-mount staining technique, utilizing the repeated application of fluorescent in situ hybridization. This technique, termed mFISH3D, was designed by extensively dissecting the chemical basis of hybridization reactions in fixed tissue. mFISH3D enabled the visualization of 10 types of mRNAs in an intact mouse brain and has been demonstrated in various biological specimens including the human brain. To achieve unprecedented levels of accuracy in spatial cell mapping, we developed artificial intelligence (AI)-driven workflow using self-supervised learning, significantly reducing the need for extensive manual annotations. The integration of mFISH3D with our AI solution sets a standard for high-dimensional tissue analysis, provides a new systematic framework for analyzing complex cellular ecosystems and enables comprehensive investigation of selective cellular vulnerabilities in diseases.

## INTRODUCTION

Understanding the function of an organ as an integrated network of distinct cell types is a central challenge in advancing medical science. Abnormal molecular events or cell loss can disrupt healthy communication between cells in a tissue, resulting in developmental defects and/or disease. Identifying such vulnerabilities with cell-type specificity is critical for formulating therapeutic strategies. However, we currently lack a research tool that enables the systematic identification of selective cell vulnerabilities across multiple cell types in a whole organ. While mRNA sequencing is a widely used tool for this purpose, its ability to quantify cell populations remains uncertain. More importantly, it does not preserve intact spatial information, which is essential for understanding how cells with different functional properties are assembled and interact.

Since all biological organisms develop through cellular proliferation, migration, differentiation, and death in an organized three-dimensional space, understanding the developmental trajectory of organs and organisms requires seamless volumetric imaging at single-cell resolution. Whole-organ clearing and imaging with light-sheet microscopy can meet this demand, and several cell-labeling techniques have been developed and integrated into this workflow. Historically, researchers have used transgenic mice with fluorescent proteins to label specific cell types for tissue clearing (Dodt et al., 2007; Ertürk et al., 2012; Chung et al., 2013; Ke et al., 2013; Susaki et al., 2014; Hama et al., 2015). A critical limitation of this approach is the time-consuming process of breeding a genetically modified strain with fluorescent-protein expression, and the necessity for genetic modification makes this approach unsuitable for the research of human tissue. While intact organ immunostaining has gradually gained popularity (Renier et al., 2014; Murray et al., 2015; Susaki et al., 2020), the slow speed of labeling (∼2 weeks for an intact mouse brain) or the complexity of the experimental setup (Mai et al., 2024; Yun et al., 2025) hinders multiplexed staining on large samples, such as a whole-mouse brain. Recently, whole-mount fluorescent in situ hybridization (FISH) has been demonstrated (Sylwestrak et al., 2016; Guo et al., 2019; Kumar et al., 2021; Kanatani et al., 2024). While this technique has not been extensively evaluated in tissues other than rodent brains, the fast speed of staining makes it an attractive option and opens the possibility for whole-mount multiplexed staining. Multiplexed staining in 2D FISH by simply iterating the staining/destaining has been performed widely in spatial transcriptomic approaches, supporting the feasibility of multi-round staining in a tissue volume.

Spatial cell mapping at the whole-organ scale calls for fast and reliable automatic cell segmentation techniques, as manual counting at the magnitude of millions of cells would be tedious and error-prone. Many studies have proposed deep-learning-based methods for automatic cell segmentation in biological images, either for general cell segmentation purposes or specifically for light-sheet images, with a convolution-based approach, such as U-Net (Ronneberger et al., 2015; Schmidt et al., 2018; Pan et al., 2019; Stringer et al., 2021; Chen et al., 2023; Luo et al., 2025) or more recently a transformer-based model architecture (Hatamizadeh et al., 2022; Hörst et al., 2024; Archit et al., 2025; Attarpour et al., 2025; Chen et al., 2025). However, applying them directly to whole-organ analysis has been challenging due to the significant variation in image quality and staining patterns across different areas of the tissue, caused by both methodological and biological heterogeneity. This variation necessitates large training datasets for sufficient generalizability, i.e., the capacity to perform well on unseen data beyond the training dataset, which could incur a prohibitive cost for the time spent in expert-level 3D annotation. Recent advances in artificial intelligence (AI) development, such as generative language models and cross-modality models (Brown et al., 2020; Radford et al., 2021), have provided new opportunities to mitigate this limitation by shifting to a data-driven approach without annotation, i.e., leveraging large amounts of unannotated data with self-supervised learning methods such as generative pretraining, masked modeling and contrastive learning (Radford and Narasimhan, 2018; Devlin et al., 2019; Chen et al., 2020; He et al., 2022). It has thus become a promising but yet-to-be-explored road map to build powerful cell segmentation models in the light-sheet microscopy domain due to their scalability and generalizability with limited annotations. Two studies have explored self-supervised learning to reduce the cost of labor-intensive manual data annotation (Achard et al., 2024; Chen et al., 2025). Achard et al. focused on a purely unsupervised method, which relies on human-derived heuristics and has not been verified in complex scenarios, such as large image datasets with significant variation in quality. Chen et al. provided valuable insights into many recent self-supervised learning methods, but their work primarily addressed semantic segmentation, which lacks the crucial instance-level information required for our biological analysis. Additionally, their label acquisition efficiency could be further improved.

We set our goal to develop a scalable and generalizable cell mapping technique for transcriptomic imaging by harnessing the power of in situ hybridization. To this end, we rigorously analyzed the chemical factors that determine the efficiency of the hybridization reaction in volumetric tissue. By finely controlling these factors and integrating a photobleaching technique, we developed mFISH3D, which enables the labeling of various RNAs at the scale of a whole-mouse brain. We demonstrated the generalizability of mFISH3D by applying the technique to various tissue types, including a whole-mouse embryo, whole-mouse kidney, whole-squid brain, and human cerebral cortex. Our rich dataset, with a variety of cytosolic staining patterns, provides an excellent resource for training deep learning models to understand the general morphology of cells. Inspired by recent advances in artificial intelligence, we developed a powerful vision transformer model, ZenCell, that is capable of accurate whole-organ analysis in 3D with limited 2D human annotation, thanks to the data-driven approach of self-supervised pretraining on a large and diverse set of regions without human annotation. By using this workflow, we obtained a precise cell map of a whole-mouse brain defined by cell-type-specific probes. Utilization of this approach following administration of NMDA receptor antagonist demonstrated its capability for revealing detailed cell specific responses in the intact mouse brain. Furthermore, we applied ZenCell to a sample from human cingulate cortex after mFISH3D, demonstrating precise 3D neuroanatomy with cell-type specificity in a large volume sample from a NIH NeuroBioBank frozen tissue collection (Freund et al., 2018). Our study demonstrates an advanced platform for exploration of tissue-wide cellular heterogeneity in three-dimensional space in multiple tissue types and offering an improved approach for quantitative studies of organ development and disease at the single-cell level.

## RESULTS

### Rational design of the protocol for whole brain in situ hybridization

Most published protocols for in situ hybridization on fixed thick-tissue volumes consisting of four major steps: delipidation, primary oligonucleotide hybridization, hybridization chain reaction (HCR) amplification, and tissue clearing (Sylwestrak et al., 2016; Guo et al., 2019; Kumar et al., 2021; Kanatani et al., 2024). Although a study of whole-brain FISH was recently published, understanding of the chemical basis underlying the binding of oligonucleotides to mRNA and their deep penetration into tissue has been limited. Characterizing the behavior of oligonucleotides within tissue is crucial for maximizing the signal-to-noise ratio (SNR) while minimizing signal loss during multiple rounds of staining to achieve highly multiplexed staining. To enhance the SNR and to ensure sufficient penetration of oligonucleotides for large biological specimens, such as a whole-mouse brain, we conducted a quantitative analysis of each reaction step to understand how modifying reaction conditions impacts staining performance.

For the delipidation step, we employed methanol, because it is one of the simplest amphiphilic molecules, and because water-free delipidation can reduce mRNA hydrolysis, which causes the degradation of mRNA. We first quantified phospholipid content by measuring phosphocholine in brain tissue treated with methanol at different temperatures. Phospholipid content was reduced by up to 82% after 3 days of incubation at temperatures above ambient (**Fig. 1A**). We then assessed SNR of 3D FISH by targeting polyadenylation sites without HCR amplification and *Gad1* mRNA with HCR amplification. The tissues after 3D FISH were cleared using a common clearing solvent, a mixture of benzyl alcohol and benzyl benzoate (BABB) after dehydration with methanol (Dent et al., 1989). A discussion of the choice of BABB over other clearing solvents is described in **Methods**. We used the hippocampal formation (HPF) for poly(dT) and the cerebellum (CB) for *Gad1* to measure SNR because of the convenience of identifying foreground signal and background noise (**Methods**). SNR decreased when brains were treated at temperatures above 60°C (**Fig. 1B** and **Fig. S1A**), and tissue autofluorescence increased at high temperatures (**Fig. S1B**). Lower temperatures (4°C or 22°C) led to strong, blob-shaped non-specific fluorescence, likely due to residual lipids (**Fig. S1A**). Therefore, we chose to proceed with delipidation at 50°C for subsequent experiments, as this temperature can offer fast delipidation while preserving mRNA signals.

**Figure 1.**
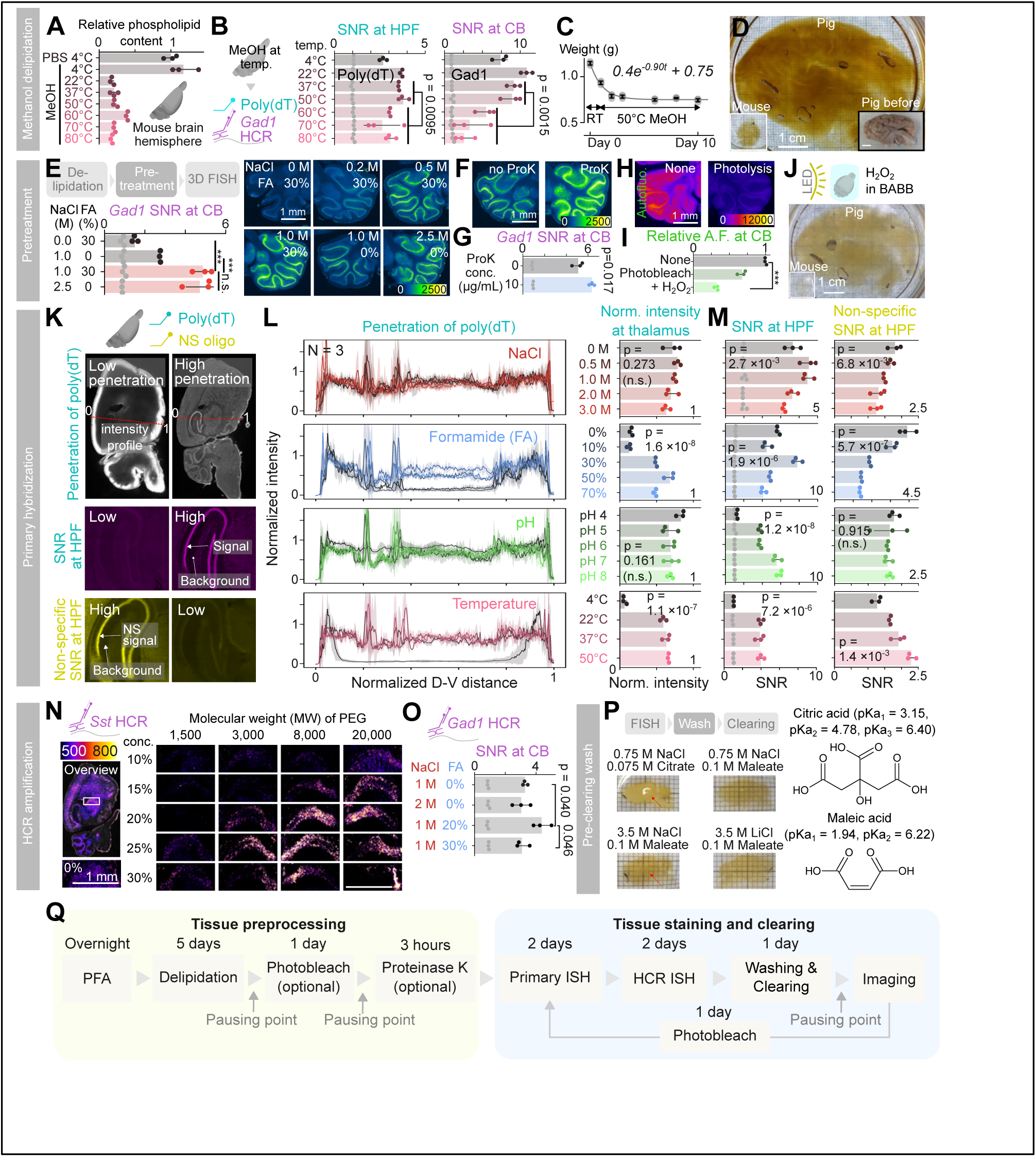
Bottom-up design of 3D fluorescent in situ hybridization. (**A**) Relative phospholipid content after delipidation in MeOH at different temperatures. The mean value of PBS control is set to one. Data are mean ± standard deviation (SD) (N = 3). (**B**) Signal-to-noise ratio (SNR) of FISH signals after MeOH delipidation at different temperatures. (Left) SNR at hippocampal formation (HFP) using poly(dT) oligonucleotides. (Right) SNR at cerebellum (CB) using HCR amplification against *Gad1*. (N = 3). Data are mean ± SD (N = 3). Wilcoxon rank sum test was used. (**C**) Weight of brain hemispheres during delipidation process at 50°C. The fitted exponential decay curve is plotted. Data are mean ± SD (N = 3). (**D**) Cleared pig brain hemisphere and whole-mouse brain are shown at the same scale. The pig brain hemisphere before clearing is shown at the bottom right corner. (**E**) Analysis of SNR of Gad1 HCR after the pre-treatment using various concentration of NaCl and formamide (FA). (Left) SNR quantification. Data are mean ± SD (N = 3). One-way ANOVA with Tukey’s post hoc test was used. *** *p* < 0.001. (Right) Representative single-z-plane images of the cerebellums. (**F**) Representative single-z-plane images of the cerebellums after Gad1 HCR with or without pretreatment of proteinase K. (**G**) Quantification of SNR of Gad1 HCR after proteinase K. Data are mean ± SD (N = 3). Student’s t-test was used. (**H**) Representative single-z-plane images of the autofluorescence in the cerebellums with or without H2O2-assisted photobleaching. (**I**) Quantification of relative autofluorescence after photobleaching. The autofluorescent intensity of the sample without photobleaching was set to one. Data are mean ± SD (N = 3). One-way ANOVA with Tukey’s post hoc test was used. *** *p* < 0.001. (**J**) Cleared pig brain hemisphere and whole-mouse brain are shown in the same scale after H2O2-assisted photobleaching. (**K**) Schematic illustration of how to evaluate the even penetration of oligonucleotide probes, SNR of specific binding, and SNR of non-specific binding. NS oligo is a fluorescent-labeled oligonucleotide to measure non-specific (NS) binding. (**L**) (Left) Intensity profile of the fluorescent-labeled poly(dT) oligonucleotides. The intensity was normalized so that the intensity at the edge became one and the intensity at the outside of the tissue is zero. Data are mean ± SD (N = 3). (Right) Quantification of the normalized intensity at thalamus. Data are mean ± SD (N = 3). One-way ANOVA was used. (**M**) Quantification of SNR of specific binding and non-specific binding in hippocampal formation. Data are mean ± SD (N = 3). One-way ANOVA was used. (**N**) Analysis of the influence of polyethylene glycol (PEG) on the penetration and amplification of HCR. Overview is shown in the left. (**O**) Quantification of the impact of salt and formamide concentration in HCR on SNR. Data are mean ± SD (N = 3). One-way ANOVA with Tukey’s post hoc test was used. (**P**) Pre-clearing washing with various buffer compositions. (Left) Photographs of cleared brains. Red arrows indicate the areas where the tissue remains opaque. (Right) chemical structure of citric acid and maleic acid. (**Q**) mFISH3D protocol. For **E**, **F**, **H**, and **N**, the intensity of the images is color-coded according to the bars.

Next, we evaluated the optimal duration for dilapidation. Brain hemisphere weight plateaued after 3 days at 50°C in methanol (**Fig. 1C**). After 3 days of incubation, HCR signals were detected in deep brain regions, such as the thalamic reticular nuclei, indicating thorough delipidation and probe penetration (**Fig. S1C**). Given that a 10-day incubation led to a 26% reduction in SNR compared to a 3-day incubation, we opted for a shorter delipidation period than 10 days (**Fig. S1D** and **Methods**). This simple delipidation protocol also enabled the clearing of larger tissues, such as an intact pig-brain hemisphere, as well as a whole-mouse brain (**Fig. 1D** and **Fig. S1E**).

Hybridization reactions typically occur in an aqueous buffer containing salt and formamide, which poses the risk of mRNA hydrolysis due to the presence of water. Hydrolysis can be accelerated by various factors, such as RNase, high pH, bivalent cations, and high temperature, but can also occur naturally in their absence.

We next explored the chemical composition of the hybridization buffer to identify a formulation that is non-destructive to mRNA. During this exploration, we found that it was difficult to determine whether signal loss was due to mRNA degradation or decreased hybridization efficiency. To independently assess the impact of buffer composition on mRNA quality, we pre-treated delipidated hemispheres with buffers of various chemical compositions before proceeding with hybridization. Hybridization was then performed using a common buffer with a fixed chemical composition. We found that pre-treatment with a low-salt buffer negatively impacted SNR (**Fig. 1E**), while adding any monovalent salt (LiCl, NaCl, CsCl) mitigated this effect (**Fig. S1F**). The absence of formamide in the buffer reduced SNR at a 1 M salt concentration, while the addition of 30% formamide further improved SNR. We hypothesized that the high salt concentration and formamide protect mRNA from RNase degradation. To test this possibility, we treated brains with RNase A in buffers with varying salt and formamide concentrations. SNR was reduced in brains treated with RNase A in buffers with low salt or formamide concentrations, but not in buffers with both high salt and high formamide concentrations (**Fig. S1G**). Based on these observations, we included either 2.5 M salt or 1.0 M salt with formamide in all subsequent reaction buffers.

Protease digestion before FISH has been known to enhance SNR of hybridization (Young et al., 2020). Proteinase K, the most common protease for FISH, typically uses a low-salt buffer without formamide. Since low salt and the absence of formamide can lead to mRNA degradation, we tested whether proteinase K could be used in a high-salt buffer with formamide. We observed a significant increase in SNR when using proteinase K with a buffer containing 1 M NaCl and 40% formamide (**Fig. 1F** and **G**).

We next sought to reduce autofluorescence through chemical pre-treatment. We had already observed that high-temperature delipidation in methanol increases autofluorescence (**Fig. S1B**). To reduce autofluorescence, we employed photobleaching because it is known to have little or no adverse effects on mRNAs and has been successfully applied to cleared tissue (Ku et al., 2020; Darche et al., 2023; Park et al., 2024). Benzyl alcohol is slightly soluble in water, which allowed us to mix hydrogen peroxide with BABB and test whether the combination of hydrogen peroxide and photobleaching in a cleared tissue could further reduce autofluorescence by accelerating the oxidation of endogenous pigments in a photobleaching device (Murakami et al., 2025). After confirming that hydrogen peroxide in BABB maintained tissue transparency, we found that autofluorescence reduction reached up to 67.9% after 16 hours of illumination with H2O2 and 32% without H2O2 (**Fig. 1H** and **I**). This H2O2-assisted photobleaching was effective in a very large tissue, such as a pig-brain hemisphere (**Fig. 1J**). Importantly, neither regular photobleaching nor H2O2-assisted photobleaching reduced FISH signal intensity (**Fig. S1H** and **S1I**). We therefore incorporated photobleaching into our standard 3D FISH workflow. We note that, as mixing hydrogen peroxide and benzyl alcohol at higher concentrations than are used here can form the potentially explosive chemical peroxybenzoic acid, we have also developed a safer alternative that does not generate an organic peroxide (**Fig. S1H, S1J**, **Methods**, and **Supplementary Note**).

Hybridization efficiency is influenced by four major factors: salt concentration, formamide concentration, pH, and temperature. While the effects of these factors on DNA-DNA hybridization are well understood (Hutton, 1977; Hillen et al., 1981), their impact on oligonucleotide penetration and binding in thick tissues is less clear. Using Cy5-conjugated poly(dT) oligonucleotides, we examined nucleotide penetration under various conditions (**Fig. 1K**, **1L** and **S1K**). Formamide concentration and temperature significantly impacted oligonucleotide penetration, while salt concentration and pH had minimal or no effect. We concluded that maintaining a formamide concentration of more than 30% and a temperature above room temperature during hybridization is crucial for penetration. We also assessed how these factors influence hybridization specificity. Ideally, hybridization should occur only when anti-sense target sites are available. Thus, to quantify non-specific binding, we designed a Cy3-conjugated oligonucleotide lacking an anti-sense binding site of more than 14 bases in the mouse transcriptome library. We analyzed both targeted binding and non-targeted binding at HPF (**Fig. 1M**). Targeted hybridization was strongest at 0.5 and 1.0 M salt, while higher salt concentrations more effectively suppressed non-specific hybridization. Interestingly, this finding contrasts with standard hybridization reactions, where higher salt concentrations typically promote more non-specific hybridization (Young et al., 2020). The reduction in non-specific binding at higher salt concentrations may be due to the neutralization of ionic attractions between oligonucleotide probes and proteins in the tissue. A formamide concentration of 30% provided the best SNR for targeted hybridization, while higher concentrations better suppressed non-specific binding. pH only affected targeted hybridization, with higher pH resulting in better SNR. We did not test pH values above 9.0, as alkaline pH can degrade mRNA and complicate data interpretation. Considering how each factor contributes to penetration, targeted hybridization, and non-specific hybridization, we chose 1 M NaCl, 40% formamide, pH 7, and 37°C as the primary hybridization conditions. These buffer conditions satisfy the condition shown in **Fig. 1E** that is required to prevent the mRNA from degradation.

We next determined the optimal duration for primary hybridization by examining how hybridization duration affects the SNR of Poly(dT)-Cy5 signals, showing that the signals plateaued after 40 hours (**Fig. S1L**). We considered the effects of inclusion of additional chemicals in the hybridization buffer. Previous investigators have found that adding chemicals such as detergents, heparin, Denhardt’s solution, and yeast tRNA can enhance the SNR of FISH in thin-tissue sections (Denhardt, 1966; Simmel et al., 2019; Young et al., 2020). We tested the effects of these chemicals and RNase inhibitor, polyvinyl sulfonic acid (Kanatani et al., 2024), in mouse brain hemispheres by mixing them into the primary hybridization buffer and found that none significantly improved SNR (**Fig. S1M**). Since these additives did not improve the reaction, they were omitted from further experiments.

We next examined the reaction solution for HCR amplification. The Pierce laboratory where HCR amplification was first developed uses a buffer made of 0.75 M NaCl, 75 mM sodium citrate (also called 5×SSC), 0.1% Tween 20, and 10% dextran sulfate (Choi et al., 2014, 2018). All published protocols for volumetric FISH use the this buffer for HCR amplification (Sylwestrak et al., 2016; Guo et al., 2019; Kumar et al., 2021; Kanatani et al., 2024). However, there is no evidence to date that this composition is the best suited for HCR in thick tissue, so we optimized the buffer composition for the HCR amplification step. During optimization, the use of dextran sulfate was problematic because it contains sodium ion, and even if it would successfully enhance the SNR by modifying the concentration of dextran sulfate, it was challenging to determine whether the improvement results from changes in salt concentration or changes in dextran sulfate itself. The function of dextran sulfate is to cause molecular crowding, which is thought to enhance the effective concentration of the HCR probes and improve their accessibility to the target (van Gijlswijk et al., 1996; Young et al., 2020). Since molecular crowding can be induced by polymers other than dextran sulfate, we tested polyethylene glycol (PEG) instead of dextran sulfate. We chose PEG because it does not contain salt, thus making the optimization process easier, and its effects on DNA-DNA hybridization are well characterized (Knowles et al., 2011). There are two important factors that contribute to molecular crowding: the molecular weight (MW) and the concentration of the polymer. We tested various MWs and concentrations of PEG and observed their impact on signal amplification and penetration (**Fig. 1N**). As suggested by previous reports (Knowles et al., 2011), signal amplification increases with higher concentrations or larger MWs. We found that using too high a concentration of PEG can result in limited penetration of the oligonucleotide, which is clearly visible in the case of 30% PEG with MW of 20,000 (PEG(20,000)). To achieve deep penetration without non-specific amplification at very high concentration of PEG, we used 10% PEG(20,000), and included formamide in the reaction solution (see **Fig. 1E**). We chose a formamide concentration of 20%, slightly reduced from 30%, because we found that 30% suppressed HCR amplification (**Fig. 1O**). We used 1 M NaCl and a temperature of 37°C with pH 7 as suggested in **Fig.1M**. We found that the penetration of HCR probes and non-specific HCR amplifications are dependent on the concentration of the primary probes (**Fig. S1N**). Importantly, this finding can explain the limited HCR probe penetration at 37°C reported by Kanatani et al. Using their TRISCO protocol, we compared high concentrations of primary probes with 4°C HCR to low concentrations with 37°C HCR. The results showed that 37°C HCR achieved uniform staining and stronger signals than 4°C HCR when a low primary probe concentration was used (**Fig. S1O**). Non-specific binding in blood vessels, commonly observed with the TRISCO protocol, was also reduced at 37°C. These findings suggest that HCR probe penetration can be enhanced without compromising signal intensity by simply lowering the concentration of primary probes.

For solvent-based tissue clearing, it is necessary to dehydrate the tissue using an amphiphilic solvent such as methanol. The use of a common sodium citrate hybridization buffer before the dehydration process resulted in an opaque tissue after the clearing process (**Fig. 1P**). This problem was reported in a previous study and solved by using tris(hydroxymethyl)aminomethane (Tris) instead of sodium citrate (Kanatani et al., 2024), but their findings did not measure the retention of mRNA after clearing. We reasoned that the opacity resulted from the limited solubility of sodium chloride and sodium citrate in methanol (**Fig. S1P**). Replacing sodium chloride with lithium chloride, and replacing citrate with maleate, which has a buffering capacity between weak acidic and neutral pH, in the buffer before the dehydration process produced tissues with high transparency (**Fig. 1P**). It is important to note that the concentration of lithium chloride can be as high as 3.5 M, which is sufficient to retain mRNA, as discussed in **Fig. 1E**. We employed 2.5 M LiCl and 0.1 M maleate buffer as the washing buffer before the dehydration process.

The sudden transition from an ionic aqueous solution to methanol caused aggregation of residual oligonucleotides, which appear as deposits in the hippocampal area. Though the signal from these deposits is faint, their exclusion can help avoid potential issues when implementing multi-cycle staining. We found that the addition of lithium chloride in methanol could remove these deposits (**Fig. S1Q**). By combining all the knowledge above, we designed our final 3D FISH protocol (**Fig. 1Q**). Using *Gad1*, a pan-inhibitory neuronal marker gene, and *Prkcd*, a thalamic regional marker gene, we confirmed the staining is even and deep enough to cover an intact mouse brain with stronger signal than TRISCO (**Fig. S1R**, **Supplementary Video 1** and **Methods**).

### Ten-plex imaging of mRNA in a whole-mouse brain using mFISH3D

In a mammalian brain, the major types of neurons are excitatory and inhibitory, and there are subclasses of neurons under each category. Identifying these subclasses of neurons is important to understand the cell-type-specific vulnerabilities in disease. Multiplexed staining of mRNAs can address this complexity. Thus, we took a simple yet powerful approach to achieve multiplexed staining in 3D tissue volumes: repetition of the staining, imaging, and de-staining processes. Using different wavelengths for excitation lasers, 488 nm, 561 nm, 640 nm, and 785 nm, one can image the spatial expression pattern of up to three types of mRNAs in addition to a counterstain. To enable registration of the images from different imaging rounds, we labeled ribosomal RNA (rRNA) as a counterstain excited at 488 nm for all rounds of the imaging. Repetition of imaging rounds linearly scaled the number of genes that we could label. This strategy has been commonly practiced in the field of spatial transcriptomics and has been demonstrated in the context of tissue clearing (Murray et al., 2015; Park et al., 2024), but has never been reported in tissue as thick as whole-mouse brain.

One main challenges in multi-round, multiplexed staining is the complete removal of the fluorescent molecules from the previous imaging round in a non-destructive way so that another set of mRNAs can be analyzed in the next round. Photobleaching perfectly fits this purpose because we can bleach the fluorescent molecules without affecting the signals in the next imaging cycle (**Fig. S1I**). Sixteen hours of H2O2-assisted photobleaching largely diminishes signals from the previous round (**Fig. S2A**). For complete removal of the fluorescent molecules from the previous round, we added 16 hours of dehybridization. Dehybridization was accomplished by incubation in 1 M NaCl and 40% formamide solution at 37°C after each round, and this condition is identical to the primary hybridization condition. We confirmed that the combination of photobleaching and dehybridization successfully removed fluorescence (**Fig. S2A**). We tested up to four rounds of staining/bleaching to demonstrate that the signals remain at the 4^th^ round with only a very minor increase in non-specific signal (**Fig. S2B and S2C**).

Another major challenge in accomplishing our goal is image analysis, particularly the registration of large 3D images with single-cell precision in the presence of both global and local deformations. While the registration of a thick tissue section has been reported in spatial transcriptomics (Wang et al., 2021), precise registration has not been attempted at the scale of a whole-mouse brain. This large scale imposes additional challenges related to global deformation, e.g. distortion of the relative distance between the cerebrum and cerebellum, as well as the local displacement, e.g. expansion or shrinkage of intracellular distances.

To develop an optimal image analysis pipeline, we prepared an rRNA-stained whole-mouse brain using our staining protocol. To obtain single-cell-resolution imaging, we set the voxel size to be 1.3 µm in the lateral direction and 2.0 µm in the axial direction, which is small enough to resolve the cells in the cleared brain despite shrinkage. The lateral optical resolution and light-sheet waist are described in **Methods**. Imaging was performed by dividing the whole brain into 4 by 4 z-stack views, and the stacks were stitched at single-cell precision using BigStitcher (**Methods**). After imaging the first round, the brain was subjected to H2O2-assisted photobleaching and dehybridization, and rRNA was stained again for the second round of imaging. The data from one channel in one round of images is 200∼300 GB. After making a series of downsampled images, also called a pyramid, we first linearly registered the lowest-resolution set of rRNA stained images with an affine transformation, followed by non-linear registration to roughly match the global shape of the images (**Fig. S2D**). To further refine the registration, we used the higher-resolution image set. Because the dataset cannot fit into the RAM of a regular workstation, we subdivided the images into multiple chunks and performed the registration in a chunk-wise manner as shown in a previous study (Wang et al., 2021). The resulting registration is highly accurate over the whole brain with single-cell-resolution precision (**Fig. S2D** and **Supplementary Video 2**).

Taking together, these experiments and image analysis resulted in an optimized workflow for multiplexed in situ hybridization in 3D (mFISH3D-2.0, and written as mFISH3D in the following). To demonstrate mFISH3D, we designed the primary oligonucleotides that specifically targets mRNAs of *Slc17a7*, *Tac1*, *Pvalb*, *Vip*, *Cck*, *Plp1*, *Calb1*, *Npy*, *Sst* and *Gad1*. We also included antibody staining using alpha-smooth muscle actin (αSMA) and a modified iDISCO staining protocol after mFISH3D. We confirmed deep whole-brain staining from mFISH3D (**Fig. 2**, **S3** and **Supplementary Video 3 and 4**).

**Figure 2.**
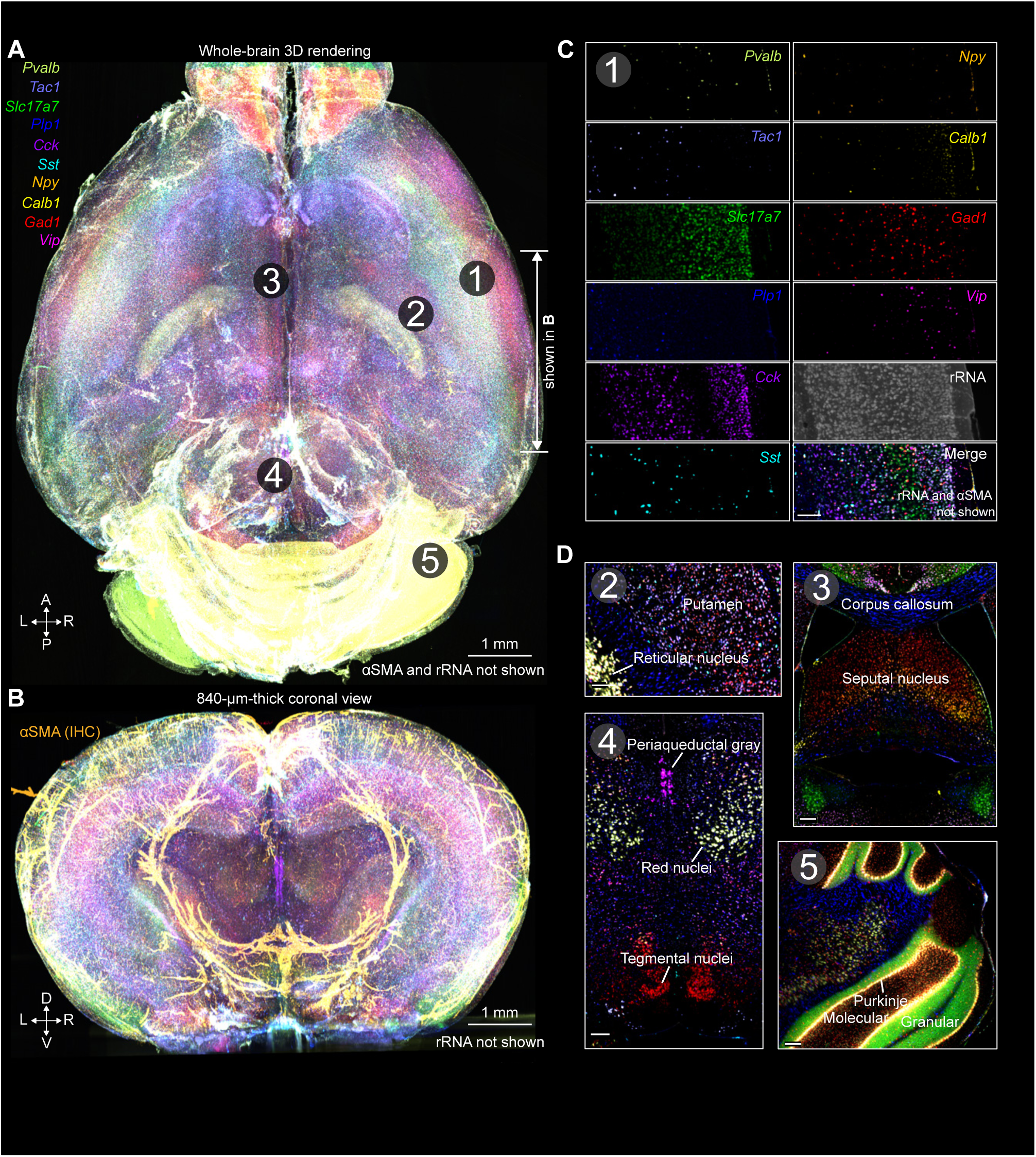
Multiplexed fluorescent in situ hybridization (mFISH3D) in a whole-mouse brain. (**A**) Whole-brain 3D rendering of mFISH3D after 4 rounds of staining. rRNA and αSMA were omitted for better visibility. L: left, R: right, A: anterior, P: posterior (**B**) Coronal view of the 3D rendering. The 840-μm-thick volume from **A** is shown. rRNA was omitted for better visibility. D: dorsal, V: ventral. (**C**) Representative magnified z-planes. The views were obtained from position 1 shown in **A**. Scale bar is 100 μm. (**D**) Representative magnified z-planes. The contrasts were adjusted depending on the areas shown and rRNA and αSMA were omitted for better visibility. The views were obtained from position 2∼5 shown in **A**. Scale bars are 100 μm.

### Application of the mFISH to various types of organs

It is a long-standing question how the expression of specific transcription factors or enzymes in a developmental pathway affects the differentiation, proliferation and migration of certain subpopulations of cells. mFISH3D holds promise of simultaneous visualization of these elements, and can further expand the realm of 3D developmental biology. We applied mFISH3D to mouse embryos (E13.5) by targeting the following four gene transcripts: *Gata2*, a zinc-finger transcription factor, whose alteration is known to be related to various diseases including hearing loss (Spinner et al., 2014), *Cyp26b1*, a monooxygenase Cytochrome P450 family member that plays an important role in tissue organogenesis as in the inner ear and spinal cord by regulating tissue retinoic acid levels (Isoherranen and Zhong, 2019), *Calb1*, a calcium-binding protein which is involved in calcium transport and/or buffering, and *Gad1*, an enzyme that catalyzes the decarboxylation of glutamate to gamma-aminobutyric acid (GABA) and is expressed in the diencephalon, ganglionic eminence (GE), and spinal cord. Along with these four transcripts, we included poly(dT) staining as a counterstain. To demonstrate the specificity of mFISH3D in mouse embryos, we included side-by-side comparisons with the developing mouse brain atlas from Allen Brain Institute (**Fig. S4**).

As shown in **Fig. 3A, 3B, 3C** and **Supplementary Video 5**, the penetration of the oligonucleotide probes in the embryo is deep enough to stain the semi-solid structures. For example, we observed even staining over a nasal turbinate, which is a curvy bony structure with a complex shape (**Fig. 3B**). We also analyzed the inner ear because it is characterized by functionally distinct components with a complex shape that includes the semi-circular canals, the vestibule, and the cochlea. Previous studies have independently reported the existence of *Gata2*, *Cyp26b1*, *Calb1* and *Gad1* in the inner ear, but an integrative understanding of the spatial information from published two-dimensional histology is challenging without a three-dimensional perspective (Meza, 2008; Buckiová and Syka, 2009; Haugas et al., 2010; Ono et al., 2020). Using mFISH3D, we can easily visualize how distinct cell populations are distributed relative to the gene products associated with development (**Fig. 3C**).

**Figure 3.**
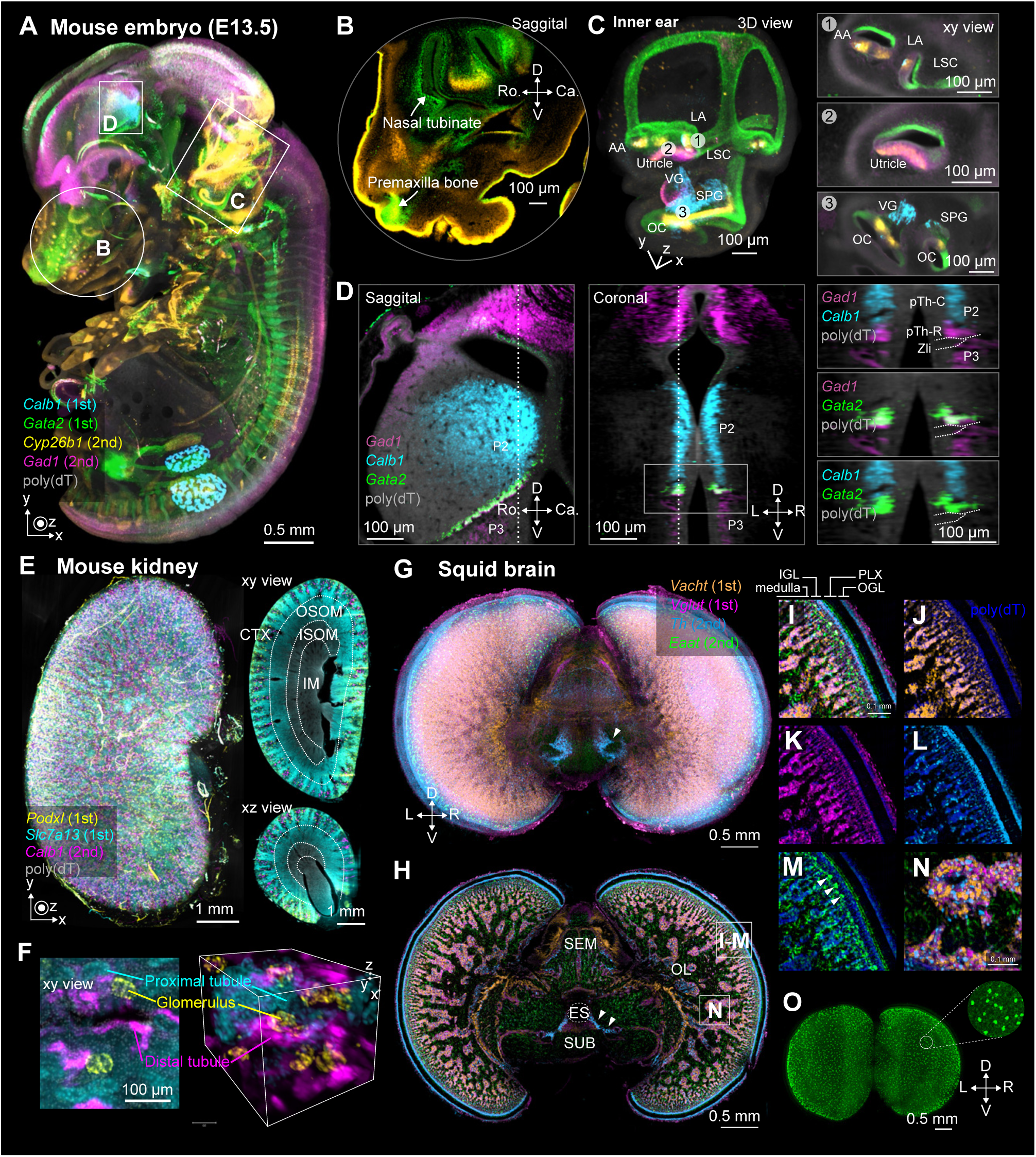
Application of mFISH3D in a mouse embryo, an adult mouse kidney and a whole-squid brain. (**A**) Volumetric rendering of whole E13.5 mouse embryo after two-round of mFISH3D. Four mRNAs are shown, and poly(dT) was used as a counter staining. (**B**) Magnified representative z-plane of the nasal cavity. D: dorsal, V: ventral, Ro.: rostral, Ca.: caudal (**C**) (Left) Volumetric rendering of the left inner ear. The other areas are not shown. AA: anterior ampulla, LA: lateral ampulla, LSC: lateral semicircular canal, VG: vestibular ganglion, SPG: spiral ganglion, OC: organ of Corti. (Right) Magnified representative z-planes of the inner ear. (**D**) (Left) Representative sagittal and coronal planes of thalamus. *Cyp26b1* is not shown due to the low expression in the area. L: left, R: right. The dotted lines indicate at which position the sagittal/coronal planes were generated. The area shown with the square solid line is shown in the right panels. (Right) Magnified coronal views of the thalamus. Up to two types of mRNAs are shown in each panel for the visualization purpose. P2: prosomere 2, P3: prosomere 3, pTh-C: caudal thalamic progenitor domain, pTh-R: rostral thalamic progenitor domain, Zli: zona limitans intrathalamica. (**E**) (Left) Volumetric rendering of an adult mouse kidney after two-round of mFISH3D. Three mRNAs are shown, and poly(dT) was used as a counter stain. (Right) Representative sectional views. CTX: cortex. OSOM: outer stripe of the outer medulla, ISOM: inner stripe of the outer medulla, IM: inner medulla. The dotted lines indicate the boundaries. (**F**) Magnified views. (Left) representative z-plane. (Right) volumetric rendering. (**G**) Volumetric rendering of a whole-squid brain after two-round of mFISH3D. Four mRNAs are shown, and poly(dT) was used as a counter staining. (**H**) Coronal section view of (**G**). The areas in white squares are shown in **I**-**M** and **N** at higher magnifications. ES: esophagus, SEM: supraesophageal mass, SUB: subesophageal mass. (**I**-**M**) A magnified view of the superficial OL, revealing sublayer structures defined by molecular expression. (**N**) A magnified view of the medulla. (**O**) A 3D rendering of *Eaat*, highlighting the large glial cell population tiling the inner surface of the IGL. IGL: inner granule cell layer, OGL: outer granule cell layer, PLX: plexiform layer.

Next, we examined whether mFISH3D can improve our understanding of differentiation of neurons in an intact brain. Though the origin of the GABAergic neurons is well characterized in the brain, little is known about the developmental origin of *Calb1* expressing cells, which is classically considered to be associated with GABAergic neurons. We found that the *Calb1*-expressing cells are predominantly localized in the dorsal thalamus (or called prosomere 2, P2) as previously reported in rat embryo (**Fig. 3D**) (Enderlin et al., 1987). Interestingly, the expression of *Calb1* did not overlap with GABAergic neurons in P2. *Gata2* is known to regulate the neurotransmitter identity of cells by promoting GABAergic and inhibiting glutamatergic neuron differentiation (Virolainen et al., 2012) and localize in the rostral thalamic progenitor domain (pTh-R) of P2. The absence of either *Gad1* or *Gata2* in *Calb1*-expressing cells suggests that the *Calb1*-expressing cells in this structure have a different cellular lineage from GABAergic neurons. Supporting this idea, *Calb1* signals were mainly found in the caudal thalamic progenitor domain (pTh-C), where glutamatergic neurons mainly exist (Virolainen et al., 2012). Further study with the inclusion of excitatory neuronal markers in mFISH3D would clarify the developmental origin of the thalamic *Calb1* positive neurons.

To demonstrate the utility of mFISH3D for the structural investigation of a non-neuronal organ, we have stained a mouse kidney against *Podxl*, *Slc7a13*, and *Calb1* (**Fig. 3E** and **Supplementary Video 6**). *Podxl* is a transmembrane protein involved in the morphogenesis of podocytes and expressed in podocytes in glomeruli (Kim et al., 2017b), *Slc7a13* is a solute carrier transporter and found in proximal tubule, and *Calb1* is found in distal convoluted tubule (Raciti et al., 2008). We used poly(dT) as a reference for registration. We stained *Podxl*, and *Slc7a13* at the first staining round and *Calb1* at the second staining round to demonstrate the compatibility with multiple rounds of staining. The signals of *Slc7a13* are visible clearly in areas as deep as the outer medulla. This is consistent with a previous report (Raciti et al., 2008) and validates the deep staining of mFISH3D in non-neuronal tissue. Together, visualization was achieved in the tertiary structure of nephrons, showing how podocytes are spaced in the proximal convoluted tubules and how tubules with distinct functional roles are positioned in 3D (**Fig. 3F**).

mFISH3D can be applied further to study the spatial gene expression of emerging model organisms. Due to a limited repertoire of antibodies available for non-mammalian species, FISH provides a faster and more scalable approach than immunohistochemistry (IHC) in building 3D gene expression brain atlases of diverse organisms (Woych et al., 2022). High-quality 3D brain atlases enable quantitative comparisons and integration across individuals, facilitating data-driven discoveries of brain structure and function. To demonstrate this concept, we studied the brain of the big fin reef squid (Sepioteuthis lessoniana). Genetically defined cell types have recently been described in areas of the cephalopod brain, utilizing single-cell RNA sequencing (Gavriouchkina et al., 2022; Songco-Casey et al., 2022; Styfhals et al., 2022; Duruz et al., 2023). However, the spatial distribution of neural cell types in their full 3D context remains largely unexplored. We designed oligonucleotide probes against *Vglut*, *Vacht*, *Th* and *Eaat* mRNAs to label glutamatergic, cholinergic, dopaminergic and glial cells, respectively, and performed mFISH3D and whole-brain imaging. mFISH3D homogeneously and brightly stained the entire squid brain despite its radically different tissue composition, including cell size and density, compared to mammalian brains (**Fig. 3G** and **3H**). The superficial optic lobe (OL) receives inputs from photoreceptor cells in the eye and is thought to be functionally analogous to mammalian retina. mFISH3D delineated the sublayer structure in this region (**Fig. 3I**). The superficial sublayer of the outer granule cell layer (OGL) was primarily glutamatergic, while the inner sublayer was enriched with dopaminergic neurons (**Fig. 3J, 3K** and **3L**). Within the plexiform layer (PLX), an inner sublayer was marked by the expression of *Eaat* mRNA (**Fig. 3M**). The inner granule cell layer (IGL) was mostly dopaminergic and glutamatergic, though an outer sublayer lacked *Vglut*-expressing cells (**Fig. 3K**). Within the OL medulla, our cellular-resolution multi-round registration pipeline revealed that almost all cells were categorized into three neurotransmitter types (glutamate, acetylcholine, and dopamine) (**Fig. 3N**), consistent with findings in other cephalopod species (Songco-Casey et al., 2022). Furthermore, at the boundary between the IGL and the medulla, we found a population of cells strongly labeled by the Eaat probe, regularly tiling the inner surface of the IGL (**Fig. 3M** and **3O**) (Gavriouchkina et al., 2022). In the supraesophageal and subesophageal mass, our probes revealed distinct expression patterns, suggesting the existence of novel brain subregions. For example, a distinct cluster of dopaminergic cells was found in the ventromedial part of the basal lobe, near the esophagus (arrowheads in **Fig. 3G and 3H**).

### AI-driven activity mapping and neuronal subtype analysis using mFISH3D

One of the important applications of mFISH3D is the quantitative mapping of cells in the brain. Despite the challenges of whole-brain-scale complexity and the absence of annotated datasets, we successfully developed robust deep learning segmentation models with minimal human intervention, leveraging recent advances in self-supervised learning. Our final model is built in a three-stage (**Fig. 4A** and **4B**) workflow to minimize annotation efforts: pre-training, 2D fine-tuning, and 3D fine-tuning. In the pre-training stage, we trained the model across all mouse brain regions using the masked autoencoding (MAE) objective, which does not require cell segmentation labels (He et al., 2022). This reduces reliance on extensive annotations and enables an efficient human-in-the-loop refinement process in later stages. The model is first fine-tuned in 2D to enable simpler initial annotations. We prepared 106 fully manually labeled 2D images of 256 × 256 pixels (333 µm × 333 µm) with 6,339 cells randomly sampled from eight image volumes targeting different genes. Subsequent annotations are obtained through proofreading of model predictions (i.e., pseudo-labels), significantly reducing human effort. **Fig 4C** presents example sets of pseudo-label obtained and manually proofread labels using an image volume targeting a gene that the model has never seen. This zero-shot prediction enabled 392 correct prediction out of 469 objects. In the final stage, 3D pseudo-labels are selected between stacked 2D predictions and native 3D predictions (**Fig. 4B**). The manually proofread 3D labels can be used to fine-tune the model to natively predict 3D segmentations skipping 2D predictions, further reducing the human effort for the manual proofreading. Our model is built on the vision transformer (ViT) architecture (Dosovitskiy et al., 2021) with the 319 million parameters with 28,480 input tokens, making it one of the largest models for cell segmentation. We also enlarge the field of view through a patch size mixing strategy, where larger patches are used to capture more background information as a trade-off for token efficiency. The large field of view has been shown to be crucial for light-sheet microscopy images, where individual object textures are inherently limited and contextual information is often decisive (**Fig. 4A** and **Methods**). Our MAE pre-training strategy greatly improved the segmentation performance on unseen test dataset as measured by the F1 score against intersection of union (IoU) threshold (**Fig. 4D**). Our final AI model, ZenCell (where “Zen” means “whole” in Japanese), achieved precise and accurate 3D cell instance segmentation, outperforming three popular baselines (StarDist, Cellpose, and Swin-UNETR) for reliable quantitative analysis (**Fig. 4D**).

**Figure 4.**
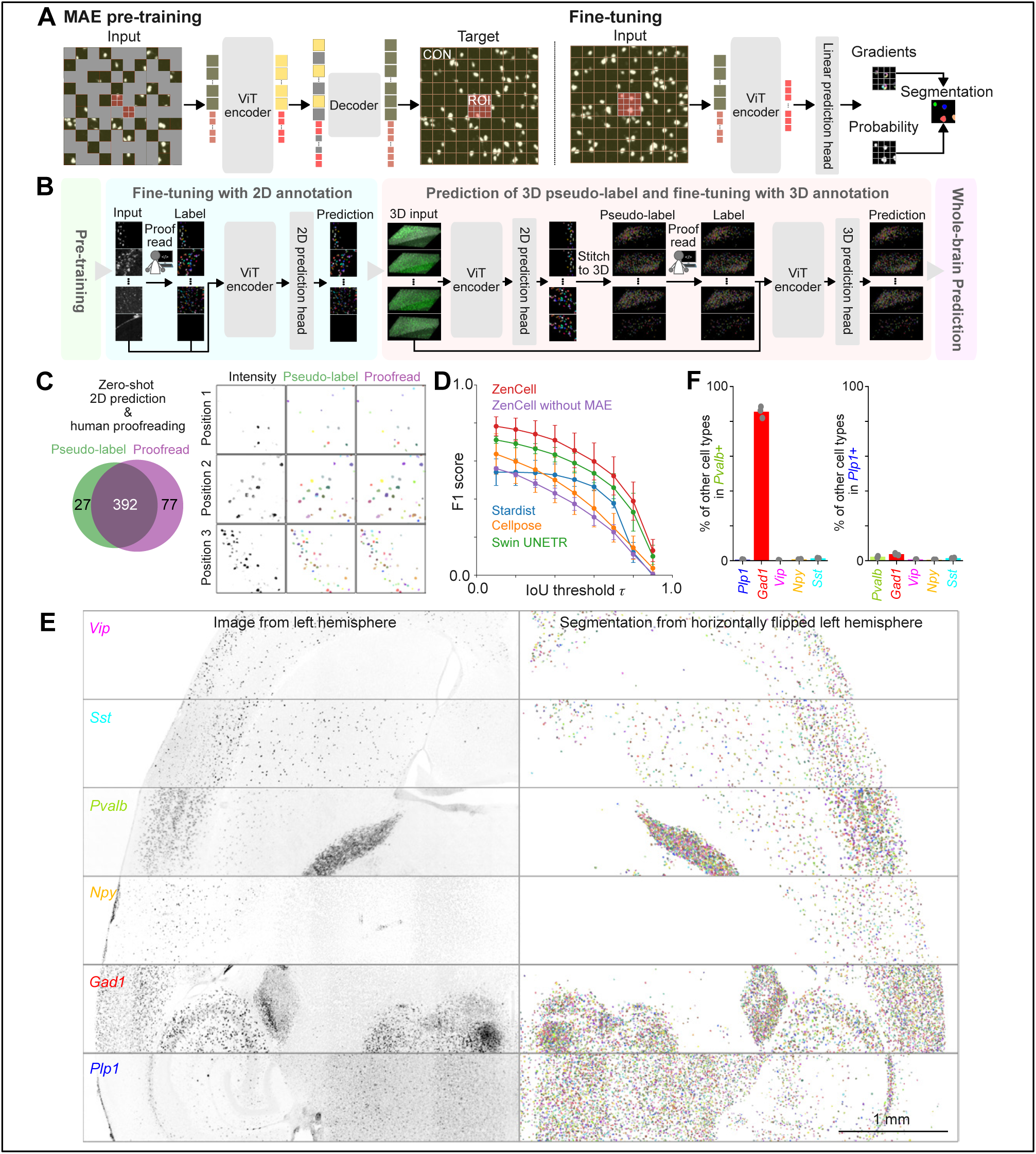
Precise whole-brain cell segmentation by leveraging context-aware pre-training of vision transformer. (**A**) Schematic illustration of the pre-training and fine-tuning process of the vision transformer. Left: context-aware pre-training with masked autoencoder (MAE). Right: fine-tuning of the pre-trained model. Right: fine-tuning of the pre-trained model. ROI: region of interest. CON: context. See Methods for details. (**B**) Schematic illustration of human annotation/correction cycles for fine-tuning of the model for whole-brain cell segmentation. (**C**) Left: Venn diagram of the number of objects identified from an unseen image of the model. Right: example images of zero-shot 2D prediction from ZenCell (Pseudo-label) and human proofread segmentations. (**D**) Comparison of cell segmentation performance across different models. The models were evaluated on unseen data across three data subsets. Data are mean ± SD. (**E**) Whole-brain segmentation using ZenCell. Left: images of 100-µm-thick z-projections from representative positions in the whole brain. Right: segmentation results of the images shown in the left. The results are horizontally flipped for visualization purposes. (**F**) Percentage of other cell types in *Pvalb*+ cells or *Plp1*+ cells in the cerebral cortex (N=3).

We performed a whole-brain analysis of cells by predicting instance segmentations for each channel using ZenCell. First, we profiled the spatial patterns of inhibitory neurons (*Gad1+*) and their subtypes (*Sst+*, *Pvalb+*, *Vip+* and *Npy+*), and oligodendrocytes (*Plp1+*) (**Fig. 4E** and **Supplementary Video 7**), demonstrating precise segmentation across the whole brain. To validate ZenCell, we performed quantitative analysis using three mouse brains. After the segmentation, we analyzed cells with multiple probes. The results showed that the percentage of *Gad1*+ cells within cortical *Pvalb*+ cells was high (∼85%), while the percentage of co-labeling with *Plp1*, *Vip*, *Npy*, and *Sst* in cortical *Pvalb*+ cells was low (less than 1.5%) (**Fig. 4F**). This agrees with the reports that *Pvalb+* interneurons are GABAergic and form a largely independent subclass from *Sst*+ and *Vip*+ interneurons with minor overlap (Rudy et al., 2011; Kim et al., 2017a). We performed the same analysis on *Plp1*+ cells, which showed minimal overlap with other cell types, consistent with the fact that *Plp1* is a marker gene for oligodendrocytes (Inamura et al., 2012) (**Fig. 4F**). To quantify the cell numbers for each anatomical annotation, we registered the brains to the Allen Brain Atlas CCFv3. Our quantification was consistent with previous quantifications in inhibitory neurons, supporting the validity of our model (**Fig. S5A**) (Kim et al., 2017a; Wang et al., 2020a). A summary of the quantification is shown in **Table S1**.

We next applied ZenCell for quantitative mapping of cellular activity following pharmacological perturbation. We designed primary probes to stain *Fos*, an immediate early gene, and performed mFISH3D on the brains of mice acutely administered the NMDA glutamate receptor antagonist, MK-801 (**Fig. 5A**). A clear distinction was observed in the patterns of *Fos* expression between the saline-administered individual (MK-, N=1) and the MK-801-administered group (MK+, N=2) (**Fig. 5B**). We also compared our expression pattern of *Fos* in MK+ mice with 3D immunohistochemistry (Susaki et al., 2020) and confirmed that the expression patterns were nearly identical with the exception of enhanced mRNA expression in the entorhinal cortex (**Fig. S5B**). While whole-brain activity mapping provides deep insights into how brain regions react to specific stimuli, multiplexed labeling of immediate early genes alongside cell-type marker genes made possible with mFISH3D provides information on both cell-type and activity patterns. In addition to *Fos* mRNA, we targeted the mRNAs of *Slc17a7* (marker gene for excitatory neurons), *Gad1*, *Pvalb*, *Sst*, *Vip*, *Npy*, and *Plp1* to characterize select subtypes of cells.

**Figure 5.**
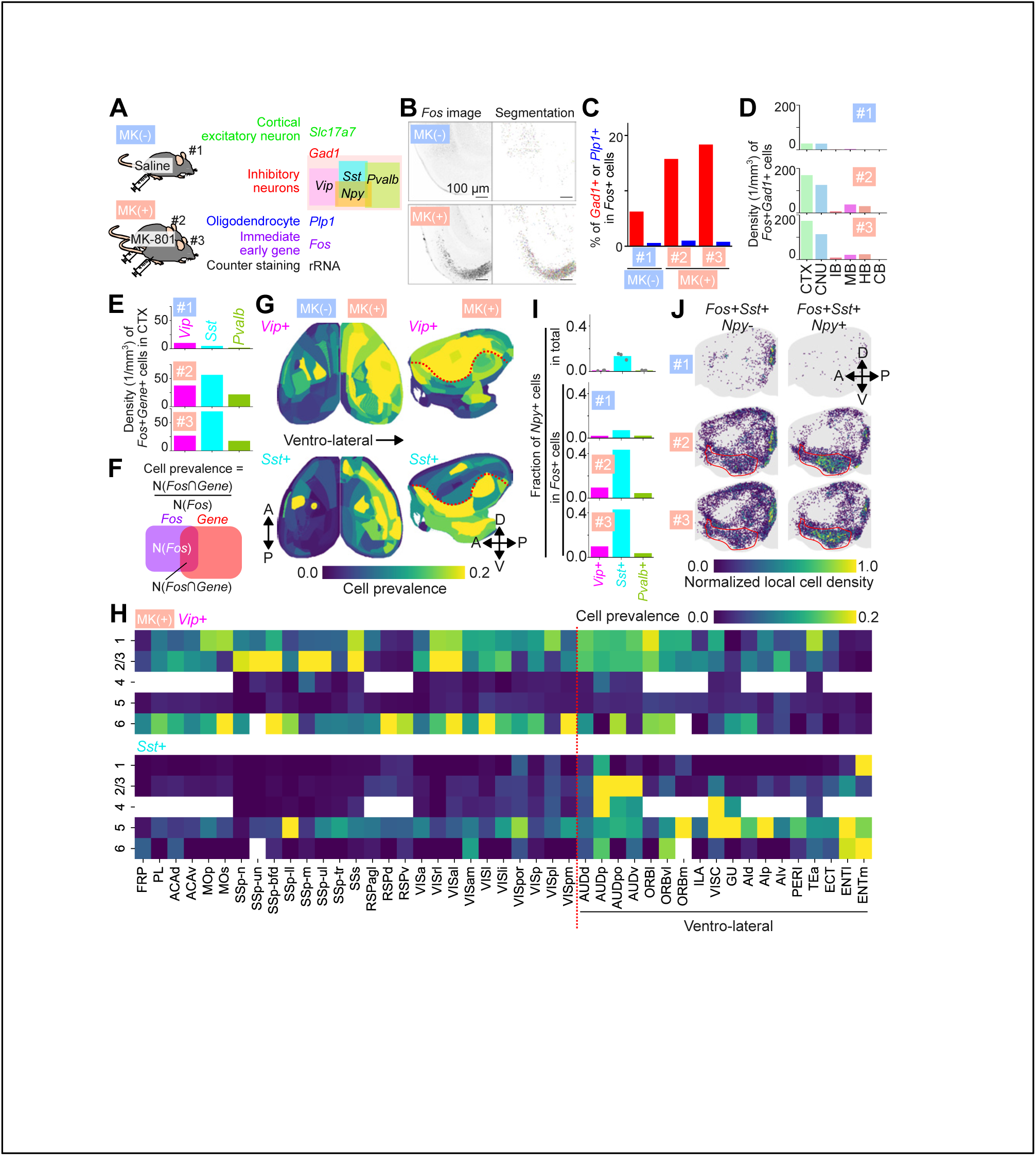
Activity mapping of inhibitory neurons after acute administration of NMDA receptor antagonist. (**A**) Illustration of the design of the experiment. The targeted mRNAs and their classifications are shown on the right. (**B**) Entorhinal cortical area of Fos-stained brains. Z-projections of 100 µm are shown for visibility purposes. (**C**) Percentages of *Gad1*+ cells (red) and *Plp1*+ cells (blue) in Fos+ cells in the brain. (**D**) Densities of *Fos*+*Gad*1+ cells in various regions of the brain. CTX: cortex, CNU: cerebral nuclei, IB: interbrain, MB: midbrain, HB: hindbrain, CB: cerebellum. (**E**) Densities of *Vip*+, *Sst*+ or *Pvalb*+ in Fos+ cells in cortex. (**F**) Calculation of cell prevalence in *Fos*+ cells. (**G**) Spatial mapping of cell prevalence in *Fos*+ cells of *Vip*+ or *Sst*+. Left: horizontal projection. Right: sagittal projection. Red dotted lines indicate the borders distinguishing ventrolateral regions from others. (**H**) Heatmap of cell prevalence in isocortex by layers. The red dotted line indicates the border distinguishing ventrolateral regions from others. (**I**) Fraction of *Npy*+ cells by cell type. (**J**) Density mapping of *Sst+Npy+* or *Sst+Npy*-cells in *Fos+* cells. Each dot indicates the position of the cells. Local cell density was calculated and shown in color. Red dotted lines indicate the piriform cortex.

We examined the subtypes of cells activated by MK-801 by quantifying the co-expression of *Fos* mRNA and cell-type marker mRNAs. It has been reported that *Fos* expression occurs in both excitatory and inhibitory neurons after administration of MK-801 (Tatsuki et al., 2016). Our analysis revealed that more than 15% of *Fos*+ cells are *Gad1*+ in MK+ mice (**Fig. 5C**). We chose to further investigate the subtype classification of *Fos*+ inhibitory neurons because *Gad1*+ cells activated upon MK-801 administration have been suggested to be associated with important biological functions, including sleep homeostasis (Tatsuki et al., 2016; Kon et al., 2024). Region analysis showed that the increase in *Fos*+ cells in inhibitory neurons is the most prominent in the cerebral cortex (**Fig. 5D**). Therefore, we focused our downstream analysis on the cerebral cortex.

One of the most basic and fundamental questions in pharmacology is the cell-type specificity of a drug. Although we have already reviewed that MK-801 can promote the activity of inhibitory neurons, additional analysis is needed to understand which subtypes of inhibitory neurons are more likely activated by MK-801. We found that the number of *Sst*+ and *Vip*+ cells are greater than *Pvalb*+ cells in *Fos*+ cells in cortex; *Sst*+*Fos*+ constituted 3.6-fold and *Vip*+*Fos*+ constituted 1.8-fold of *Pvalb*+*Fos*+ cells (**Fig. 5E**). This is intriguing given that the number of *Pvalb*+ cells is approximately equal to the number of *Sst*+ (∼1.0 fold of *Pvalb*+) or *Vip*+ neurons (∼0.59 fold of *Pvalb*+) in the cortex (**Fig. S5C**). We calculated the prevalence of a specific cell types within the *Fos*+ cells in different regions of the brain (**Fig. 5F**). There are distinctions in the spatial patterns of how *Vip*+ and *Sst*+ cells respond to the drug, depending on the region of the brain. The dorsal regions are more dominated by *Fos*+*Vip*+ cells while ventrolateral regions are more dominated by *Fos*+*Sst*+ cells (**Fig. 5G** and **S5D**). Specifically, the cell prevalence of *Fos*+*Vip*+ cells is 1.2-fold higher in dorsal regions compared to ventrolateral regions, while the cell prevalence of *Fos*+*Sst*+ cells is 3.3-fold higher in ventrolateral regions compared to dorsal regions. The dorsal part of the cortex is comprised of isocortex, which is characterized by its laminar structure. We profiled the ratio of *Fos+Vip+* or *Fos+Sst+* cells within *Fos*+ cells across each cortical region along the laminae. *Vip*+ cells are widely activated across cortical layers 1, 2, 3, and 6, while *Fos*+*Sst*+ cells are localized in layers 5 and 6 of the ventrolateral region of the isocortex (**Fig. 5H** and **S5E**).

Since the *Sst*+ cells are the most abundant inhibitory neuron types in the *Fos*+ cell population in the cortex (**Fig. 5E**), we investigated the signature of *Sst*+ cells activated by MK-801. Our data suggest that ∼13.5% *Sst*+ cells express *Npy*, but this number goes up to ∼40% if we limit our analysis to *Fos*+ cells in MK+ mice (**Fig. 5I**). We then asked whether there are any differences in spatial patterns between *Sst*+*Npy*+ and *Sst*+*Npy*-cells in response to MK-801. Our analysis revealed a particularly high density of activated *Sst*+*Npy*+ cells in the piriform cortex (**Fig. 5J**). The fraction of *Sst*+*Npy*+ cells in *Fos*+ cells was ∼75% in the piriform cortex of MK+ mice. Given the significant number of *Fos*+ cells in the piriform cortex along with entorhinal cortex, retrosplenial cortex and olfactory bulb (**Fig. S5F**), these data suggest that *Sst*+*Npy*+ inhibitory neurons have a functional role distinct from other *Sst*+ cells in response to MK-801. In summary, the drug MK-801 causes a cell-type dependent activation pattern, and spatial patterns exist within this cell-type selectivity. The broader implication of our findings is that our pipeline can quantify the cell-type and brain region dependent response of pharmaceutical therapeutics, which is crucial in drug development for assessing the mechanisms of action and the specificity of a drug.

### mFISH3D on the human brain

Since Cajal’s pioneering work in precise cell mapping, neuroscientists have devoted tremendous effort toward understanding the cellular architecture of human brain. Recent advances in transferring whole-mount staining and clearing techniques to the human brain have generated a new field of three-dimensional, cellular-resolution human brain imaging (Zhao et al., 2020; Park et al., 2024). However, technical challenges remain that are either absent or negligible in rodent brains. One obstacle is the high level of autofluorescence, particularly fluorescence from lipofuscin, which often masks true signals of interest. Using H2O2-assisted photobleaching, this problem was minimized in human tissue (Murakami et al., 2025). Another challenge is the variable quality of tissue samples available from publicly supported brain tissue banks. This variability is unavoidable due to differences in postmortem interval and the time required to cool the brains to low temperatures that cannot be fully controlled during the process of obtaining donor samples. In our experience, these considerations result in a significant number of samples from various brain banks being identified as less favorable once analyzed, despite the many precautions taken in routine acquisition of the samples. To determine whether these factors limit the utility of mFISH3D for archived human donor tissues, we prepared samples of fresh-frozen cerebral cortex from six independent donors from the Miami Endowment Brain Bank that is affiliated with NIH NeuroBrainBank. Brodmann area 9 (BA9) was used to try to ensure the reproducibility of the dissections.

We first sub-dissected the frozen block of the brain into pieces with ∼5 mm size in one dimension. We followed the mFISH3D protocol after fixation and H2O2-assisted photobleaching of autofluorescence. We stained *Npy*, *Sst* and *Plp1* and observed signals of FISH with high contrast in all six samples. These data confirming the routine utility of mFISH3D in studying mRNAs of postmortem human brains. We confirmed the specificity of the staining by verifying that staining patterns matched the known gene expression patterns: major overlaps between *Npy* and *Sst* in a subtype of interneurons and *Plp1* in oligodendrocytes (**Fig. 6A**). Lastly, we examined whether a human brain is amenable to the multiple rounds of staining using a large piece from a brain. We prepared a cingulate cortex sample (BA24) of approximately 20 mm × 10 mm × 5 mm, and performed mFISH3D against rRNA, poly(dT), *Npy*, *Sst*, *Plp1*, *Slc17a7, Cck* with IHC of αSMA (**Fig. 6B** and **Supplementary Videos 8**). The staining pattern reproduces the general cortical laminar pattern (**Fig. 6C**). To provide more interpretable data from the 3D image, we employed ZenCell to extract the positions of *Sst*+ and *Npy*+ cells. The laminar distribution of *Sst*+ cells is well characterized (Banovac et al., 2022). Quantification of *Sst*+ cell density was consistent with the report, taking into account tissue shrinkage by a factor of 2.0-2.5, revealing a higher density in layers II and IV than in other layers (**Fig. S6A** and **S6B**). The analysis of the pattern of the *Npy*+ ratio in *Sst*+ cells was also consistent with the report, with the white matter and layer VI showing higher ratios than in other layers, thus validating our AI-driven quantification (**Fig. S6C** and **S6D**). Importantly, the analysis can be expanded to the entire imaged volume, demonstrating the utility of this approach (**Fig. 6D**). Although further analysis of human samples from unaffected controls and patients suffering from diseases or developmental disorders will be required to definitively demonstrate the utility of mFISH3D in human disorders, these data show that this new approach, combining an optimized experimental protocol and AI, can be particularly powerful for studies of a variety of human medical problems.

**Figure 6.**
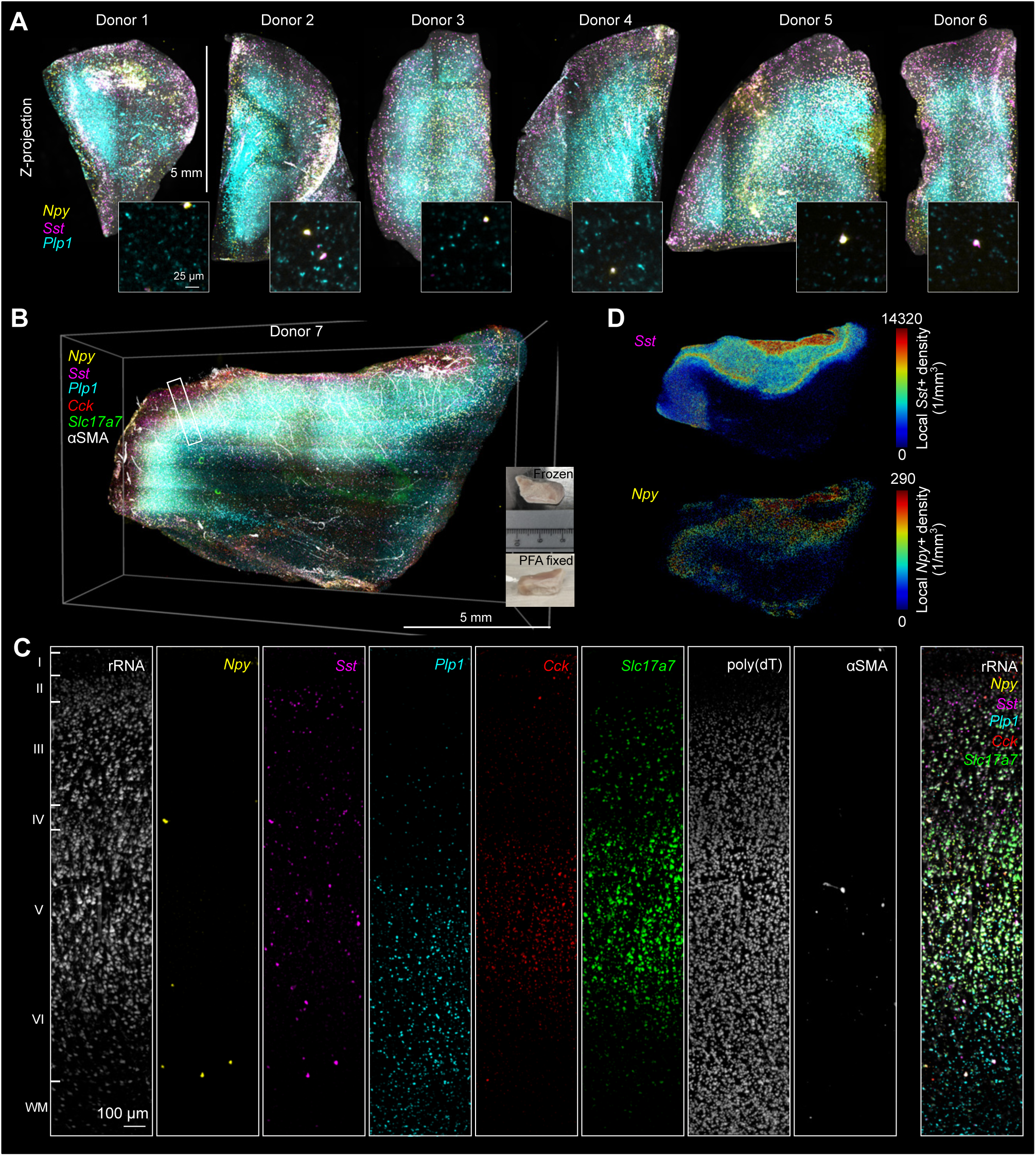
mFISH3D and ZenCell in human brain. (**A**) Application of single-round mFISH3D in a human brain from different donors. In the volume rendering, a maximum filter was applied, and the contrast was adjusted for each volume for better visibility. Magnified views of representative z-planes are shown in the insets. (**B**) Volumetric rendering of a large section from the cingulate cortex after mFISH3D. Photographs before staining are shown at the bottom right. (**C**) Magnified views of the cortical laminae. Images of 50-µm-thick z-projections from representative positions are shown. The positions of the laminae were inferred from the images and are indicated on the left. WM: white matter. (**D**) Whole-mount analysis of cells using ZenCell against *Sst* and *Npy*. Each dot indicates the position of the cell and the color indicates the local cell density.

## DISCUSSION

Analysis of gene expression and cell type abundance in large, 3D tissue volumes is a new area of investigation that has become a focus of modern neuroanatomy. We report here three important advances that can accelerate progress in developmental biology and disease: we first establish a chemically optimized protocol for delipidation, probe penetration and RNA hybridization that can be used for 3D analysis of a variety of tissue types from multiple organs and experimental systems; we then optimize photobleaching to enable multiplex, serial experimentation for detection of at least ten individual probes in large volume tissues; and we introduce an AI-driven framework for whole brain cell analysis that can be extended to incorporate additional pre-training datasets with varying resolutions, staining methods, and tissue types to enhance the model’s generalizability.

mFISH3D is based on rigorous chemical analysis of 3D FISH, followed by simplification of the protocol. By dissecting the 3D FISH reaction, we revealed chemical components necessary to protect mRNA in tissue, allowing for whole-organ 3D FISH. Our findings indicate that this bottom-up approach was important for expanding the utility of 3D FISH, enabling us to perform 3D FISH in various types of tissues, including the postmortem human brain. The chemical insights obtained in this study and simple molecular composition of mFISH3D provides a foundation for extending the technology to other domains. Extending the tissue thickness usable in FISH-based spatial transcriptomics, for example, is an exciting future direction. It seems evident that aggressive delipidation with MeOH at high temperature will facilitate deep penetration of key probes into tissues, RNA-friendly hybridization reactions will protect the RNAs from degradation within the tissue during the process, and H2O2-assisted bleaching will efficiently promote multi-round imaging.

We believe the ability to simply alter the mFISH3D protocol is an important strength of the methodology that can enable its further development for specific applications. For example, we did not include chemical modifications to the tissue using hydrogel tissue chemistry, but it seems likely that the addition of the anchoring matrices is promising for the protection of fluorescent proteins and longer retention of mRNAs (Chen et al., 2016; Ku et al., 2020; Sylwestrak et al., 2016). We note that this strategy can be easily integrated into mFISH3D by introducing gel embedding as a preprocessing step.

We have demonstrated the combination of immunostaining and mFISH3D. Although we have not tested it, there is immense potential in the use of HCR to amplify the signal from antibodies, thereby assisting simultaneous imaging of RNA and protein. The combination of HCR and immunostaining has been reported to enhance signal amplification and could provide a promising alternative to conventional antibody-based signal amplification (Lin et al., 2018). We note, however, that some antibodies are not compatible with the alcohol pre-treatment used in mFISH3D. A list of antibodies incompatible with alcohol pre-treatment can be found on (https://idisco.info/validated-antibodies/) for pre-screening antibodies of interest.

A current limitation of mFISH3D is that the maximum number of genes that can be amplified is limited by the repertoire of HCR probes. This limitation arises from the mild de-hybridization process of mFISH3D, which does not support the extensive removal of oligonucleotides from previous staining rounds. Consequently, expanding the repertoire of HCR probes could significantly enhance the potential of mFISH3D and enable even more complex staining and imaging protocols. For example, a major vendor of HCR probes, Molecular Instruments, sells 10 types of orthogonal HCR probe sets, and other studies have reported additional repertoires (Tsuneoka and Funato, 2020; Wang et al., 2020b). As HCR is a key component of the 3D FISH reaction, integrating custom-made HCR probes would add more flexibility to the signal amplification strategy in mFISH3D.

Finally, we have presented ZenCell, an AI-driven framework for whole-brain cell-type-specific analysis. ZenCell is based on pre-training using our internal datasets to recognize various cell morphologies. As progress continues, this approach can extended to incorporate a variety of pre-training datasets for even stronger generalizability. A current limitation of ZenCell is that the 3D pseudo-labeling is generated by simple vertically stacking of 2D segmentations, which leads to erroneous 3D pseudo-labels and requires manual correction. For this reason, we avoided fine-tuning in areas with a high density of neurons, such as striatal medium spiny neurons or cerebellar stellate cells, where the manual correction of 3D pseudo-label can be time-consuming. We expect that integrating multi-directional fusion of 2D segmentations will improve the quality of pseudo-labeling and significantly reduce the need for manual corrections in 3D (Zhou et al., 2024).

Ultimately, it is our goal to apply this combination of AI-driven mapping of cells and multiplexed, single-cell-resolution, whole-mount imaging to study the complex cellular ecosystem in intact human-organ systems, accelerating our understanding of development and selective cell vulnerabilities in disease and aging. Our demonstration that tissue clearing of a pig brain hemisphere can be achieved using this protocol is an initial step in this direction.

## MODEL SYSTEMS AND PERMISSIONS

### Mice Model

C57BL/6J mice from Jackson laboratory were bred by Hatten lab in The Rockefeller University and used for the experiments. All procedures involving mice were approved by The Rockefeller University Institutional Animal Care and Use Committee (IACUC) and were in accordance with the National Institutes of Health guidelines.

#### Collection of brain and kidney

The 8-week-old mice were sacrificed by an overdose of pentobarbital (> 100 mg/kg), then transcardially perfused with 15 ml of phosphate buffer saline (PBS) and 20 ml of 4% paraformaldehyde (PFA, Electron Microscopy) in PBS. The brains or kidneys were dissected, then post-fixed with 40 mL of 4% PFA in PBS at 4°C overnight. We used approximately 6 times of the volume of PFA solution to the volume of the tissue. The fixed organs were then brought to the delipidation process, which is shown below in the *Delipidation* section.

#### Collection of mouse embryo

Pregnant female mice were euthanized at embryonic day 13.5 (E13.5) by using CO₂ asphyxiation. The embryos were then dissected from the uterine sacs, and they were immediately subjected to the fixation in 40 mL of 4% PFA in PBS at 4°C overnight. The following treatment is the same as for the brains described above.

#### Drug administration

We used 8-week-old mice. We intraperitoneally administered 20 mg/kg MK-801 in saline (Sigma-Aldrich #M107) or an equal volume of saline at Zeitgeber Time (ZT) 2 and sacrificed the animals at ZT 4. We followed the procedure for brain dissection as described above.

### Squid

Squids (Sepiteuthis lessoniana) were cultured at the OIST Marine Science Station. Juvenile squids (approximately 2 cm mantle length) were euthanized by 5% ethanol in seawater. Subsequently, the brains were dissected and fixed with 4% PFA in PBS at 4°C overnight. All procedures involving squid were conducted in accordance with institutional guidelines, approved by the OIST Animal Care and Use Committee. The treatment after fixation is the same as for a mouse brain described above.

### Pig

A fresh pig brain was obtained from a local butcher and dissected into halves. We fixed the tissue with 200 mL of 4% PFA in PBS at 4°C overnight. The fixed tissue was brought to the delipidation process as in a mouse brain.

### Human Tissue

Fresh frozen post-mortem human tissue samples were obtained from The University of Miami Brain Endowment Bank. Project approval was sought and granted by The Rockefeller University Institutional Review Board. Tissue donors were de-identified before receipt of tissue. Donor metadata are described in **Table S2**. In short, for **Fig. 6A** and **6B**, the donors with PMI being less than 30 hours, and with RIN more than 5 were chosen. The age of the donors was between 65-86 years old. The donors were all male and were either control without Alzheimer’s disease or with Alzheimer’s disease Braak stage III-IV. The tissues were obtained from Brodmann area 9. For **Fig. 6C**, the 25-years-old male donor with PMI 19.4 hours was used. The tissue was obtained from Brodmann area 24. The fresh-frozen sample blocks underwent further sub-dissection. The sample blocks were taken from -80°C and transferred to -40°C for two hours. The blocks were then placed on a metal plate over dry ice and subjected for the sub-dissection. After the sub-dissection, we moved the samples to -20°C and incubated for more than 3 hours. We then fixed the tissue with 40 mL of 4% PFA in PBS at 4°C overnight. The fixed tissue was brought to the delipidation process as in a mouse brain.

## METHOD DETAILS

### mFISH3D

The most updated version of the mFISH3D protocol can be found at protocol.io (DOI: dx.doi.org/10.17504/protocols.io.kqdg3qn21v25/v1). The sequence of oligonucleotides used in the study is shown in **Table S3** and the choice of oligonucleotides used for each figure was shown in **Table S4**. The exact staining process used in this study is described below.

#### Design of oligonucleotides

We used in situ hybridization with signal amplification by hybridization chain reaction, HCR (Choi et al., 2018). The original HCR studies have provided sets of hairpin-shaped oligonucleotides (hereafter HCR probes) that have little to no mutual crosstalk. The sets of the sequences were named B1 to B10, and the sequences of the B1 to B5 were publicly available from the previous study (Choi et al., 2014) while the sequences of B6 to B10 were not disclosed (Schulte et al., 2024) and the oligonucleotides were only available through Molecular Instruments, Inc. There is another study published about HCR (Wang et al., 2020b), where the authors also described the set of orthogonal HCR probes to expand the variation of the sequences, and their sequences were publicly available. Confusingly, the expanded sequences ware also named B6, B7, …, which is unrelated to the B6, B7, … from Molecular Instruments, Inc. To avoid the confusion, we rename the sequences from Wang et al., as BW6, BW7, …. Unless we use the B6 ∼ B10 HCR probes, all the initiator-tagged DNA oligonucleotides (hereafter primary probes) are synthesized by referring to the design of the third-generation HCR (Choi et al., 2018) at Integrated DNA Technologies (IDT) otherwise purchased from Molecular Instruments, Inc. The design of the primary probe is described in the following.

We obtained the mRNA sequence from the NCBI website (https://www.ncbi.nlm.nih.gov) and generated its reverse complement. The sequence was then divided into 20∼25-mer oligonucleotides using OligoMiner (Beliveau et al., 2018), ensuring that each oligonucleotide met the criteria of at least a 1-mer gap between adjacent oligonucleotides and a GC content between 25% and 75%. We did not consider the Tm value of the oligonucleotides because the split design of the primary probes helps to reduce off-target signal amplification (Wang et al., 2012). To evaluate off-target binding, the oligonucleotides were aligned against the transcriptome database of the species of interest, which was downloaded from Ensembl (http://useast.ensembl.org/info/data/ftp/index.html). BLAST was used for the alignment. To minimize off-target signal amplification, oligonucleotides with more than 14-mer off-target binding were subjected to additional analysis. Oligonucleotides were excluded from consideration if any combination of two off-target binding sites was found in proximity of less than 100-bases threshold. The remaining oligonucleotides were tagged with sequences for HCR amplification with two spacer nucleotides. We mixed the 1 mM of each oligonucleotide at equal volume and labeled the concentration of the mixture as 1 mM primary probes. The fluorescent-dye-conjugated HCR probes were either purchased from Molecular Instruments or synthesized by IDT (Wang et al., 2020). The scripts used for designing the primary probes are available on GitHub (https://github.com/tatz-murakami/split-oligo-designer).

#### Delipidation

Unless noted, the volume of the solution for delipidation process was same amount as PFA solution used for fixation. After PFA-fixation, the fixed tissues were directly moved into 80%(v/v) MeOH solution in water at 4°C and were gently shaken for 2 hours for dehydration purposes. We skipped PBS washing, which is usually done after PFA fixation, to simplify the workflow. We confirmed that both phosphate ions and paraformaldehyde are soluble in 80% MeOH and expected the molecules to be washed out in 80% MeOH. The reason we chose 4°C is because the incubation of the tissues at high-temperature in the low-salt solution can degrade mRNA as implied in **Fig. 1E**. We repeated the dehydration using 80%(v/v) MeOH with shaking for 2 hours at 4°C. The samples were transferred to 100% MeOH with shaking for 2 hours at 4°C. After refreshing MeOH, the samples were transferred to room temperature (RT) and kept being shaken overnight. We refreshed the MeOH and transferred the sampled to 50°C with shaking for 5 days. We refreshed MeOH every day but skipped refreshments during a weekend or national holidays. If necessary, the delipidated tissues were stored at -20°C until use.

#### H2O2-assisted photobleaching of autofluorescence

For the tissues shown in **Fig. 2, 4, and 5**, we performed H2O2-assisted photobleaching (Murakami et al., 2025). Though optional, photobleaching will significantly improve the SNR (**Fig. 1I**). Note that mixing benzyl alcohol and hydrogen peroxide can assist formation of peroxybenzoic acid that may become explosive when concentrated. While we do not expect distillation to occur in the benzyl alcohol-benzyl benzoate mixture (BABB) due to the nonvolatile nature of benzyl alcohol (boiling point: 205 °C) and benzyl benzoate (boiling point: 323 °C), we do not exclude the possibility that an improper use of the reagents can lead to an accident. Therefore, we strongly recommend using benzyl acetate instead of benzyl alcohol. We first prepared BABB by mixing benzyl alcohol and benzyl benzoate at the volume ratio of 1:2. We added H2O2 to BABB so that the final concentration is 0.2%. For the safer alternative, benzyl acetate was mixed with 0.4% H2O2. This hydrogen peroxide–benzyl acetate solution was then combined with an equal volume of benzyl benzoate to produce hydrogen peroxide BAcBB so that the final concentration of H2O2 is 0.2%. Because the clearing performance of BAcBB is not as high as that of BABB, we used BAcBB only for photobleaching purposes.

Throughout this manuscript, we used BABB unless otherwise noted. However, we recommend using BAcBB due to the safety considerations described above. The solution of delipidated tissues stored in MeOH were replaced with hydrogen peroxide clearing solvent. We used 2.5 ml of the solution for a whole brain or 2.0 ml of the solution for a hemisphere brain or a smaller sample. After 1 hour of shaking, we refreshed the solution and kept shaking the sample until fully cleared. The samples were transferred to a non-colored glass vial and placed in the photobleacher. The bleaching process was performed overnight. The samples were then washed with 100% MeOH for 2 hours. We repeated the MeOH washing three times in total. If necessary, the bleached tissues were stored at -20°C until use in MeOH.

#### Primary hybridization

For this section, 1.5 mL of the solutions were used for a whole-mouse brain, or 1.0 mL for a hemisphere or the tissues with smaller volume. The tissues in MeOH were transferred to the solution with 2.5 M LiCl in 100% MeOH (hereby 2.5 M LiCl MeOH). Note the solution contains no water, and the purpose is to retain the salt concentration in a tissue to be high during the transitioning of MeOH-based solution to aqueous solution. The tissues were gently shaken in 2.5 M LiCl MeOH for 1 hour at RT. We next proceeded to the optional proteinase K treatment. For **Fig. 2 and 4**, we used proteinase K while we skipped proteinase K in **Fig. 5** for the following reason. Though the proteinase K helps to enhance SNR, the over-digestion can in turn deteriorate SNR. The time point when the over-digestion starts varying depending on the size of the tissue. As the size of the human brain pieces are not easily controllable, we excluded the proteinase K to omit the possibility that proteinase K changes the interpretation of the intensity of signals. We used primary hybridization buffer (1 M NaCl, 40% formamide in 100 mM lithium maleate buffer (pH 7.0)) for proteinase K. Lithium maleate buffer was made by mixing maleic acid and lithium hydroxide. For the digestion, the 10 µg/mL or proteinase K was added in the primary hybridization buffer, and the tissue was gently shaken at RT for 3 hours. We used 1.5 mL of the solution for a whole-mouse brain or 1.0 mL for a hemisphere. After the treatment with proteinase K, the tissue was washed with primary hybridization buffer twice by gentle shaking for 30 minutes each at RT. The primary hybridization was performed by adding primary probes in the primary hybridization buffer and gently shaking the sample at 37°C for one night (mouse hemisphere or smaller) or two nights (whole-mouse brain). Depending on the weekend schedule, we extended the process to three nights but no more than four nights. From our observation, there is no clear distinction in SNR between two nights and three nights. The concentration of the primary probes was adjusted to achieve the even staining (**Fig. S1N**). We began primary hybridization using 4.0 nM for each primary oligonucleotide, and reduced the concentration till even staining was obtained. The tissue was washed with pre-HCR washing buffer (1 M NaCl and 20% formamide in 100 mM lithium maleate buffer (pH 7.0)) with gently shaking for 2 hours three times before the HCR amplification.

#### HCR amplification

For HCR amplification, 900 µL of the solutions were used for a whole-mouse brain, or 600 µL of the solutions were used for a hemisphere or the tissues with smaller volume, and 1.5 mL or 1.0 mL were used for post-HCR washing. Tissues were directly moved to our HCR buffer (1 M NaCl, 20% formamide, and 10% PEG(20000) in 100 mM lithium maleate buffer (pH ∼7.0)) with 30∼60 nM of HCR probes. The HCR probes were denatured prior to the reaction by heating the probes to 95°C and left at RT for at least 30 minutes. The HCR amplification was performed at 37°C with a gentle shake for two nights. Depending on the weekend schedule, we extended the duration to three nights but no more than four nights. The samples were washed with post-HCR washing buffer (1 M LiCl, 40% formamide in 100 mM lithium maleate buffer (pH ∼7.0)) at 37C with gently shake for 2 hours three times.

#### Pre-clearing process and tissue clearing

For this step, 1.5 mL of the solutions were used for a whole-mouse brain, or 1.0 mL for a hemisphere or the tissues with smaller volume unless STP 120 tissue processor was used. To automate the process, we used STP 120 tissue processor filled with 1 L of the solutions. The use of the tissue processor is completely optional and intended to save time for the clearing process otherwise lengthy. The tissues were held in histology cassettes when the tissue processor was used. The tissue was moved to pre-clearing washing buffer (2.5 M LiCl in 100 mM lithium maleate buffer (pH ∼7.0)) to remove the residual NaCl and formamide. The tissue was gently shaken at RT overnight for manual processing, or 4 hours twice for tissue processor. The tissues were immersed in 2.5 M LiCl MeOH solution with gently shake for 30 min, then immersed in 100% MeOH. The tissues were shaken at RT for 30 min, and we repeated the process three times before bringing the tissues in BABB. BABB solution was refreshed every one hour until the tissue got fully transparent.

#### Photobleaching of imaged samples

We followed the same process as shown in the “H2O2-assisted photobleaching of autofluorescence” section.

#### Reason for choosing BABB

We chose BABB over other common clearing solvents such as dibenzyl ether (DBE) because of its solubility in water. BABB, or more specifically benzyl alcohol, is slightly soluble in water due to its high polarity compared to other clearing solvents. Since mFISH3D frequently switches the environment of the solution between aqueous and lipophilic, there is a possibility that the remaining chemical, e.g. water, salt, clearing solvent, could affect the next reaction. For example, the remaining water may cause the tissue to remain opaque after clearing, which has been reported in DBE (Muntifering et al., 2018). The residual clearing solvent may affect the hybridization reaction. The choice of BABB greatly reduces the risk of these events. We did not choose hydrophilic clearing reagents such as CUBIC (Matsumoto et al., 2019), because none of the aqueous clearing solutions can meet the condition in **Fig. 1E**.

#### Comparison of TRISCO and mFISH3D

For a fair comparison between TRISCO and mFISH3D, we introduced several modifications to TRISCO for the consistency. For TRISCO, reaction volumes of 1.0 mL and 600 µL were used for primary hybridization and HCR, respectively. We employed 60 nM hairpin HCR probes conjugated with Alexa Fluor 546. BABB was used for tissue clearing instead of dichloromethane and dibenzyl ether. All other steps followed the original TRISCO protocol. For mFISH3D, we followed the whole-brain protocol as described above with photobleaching and without proteinase K pretreatment. We employed 60 nM hairpin HCR probes conjugated with Alexa Fluor 546.

### Microscopy

#### Light-sheet Fluorescence Microscopy

The Cleared Tissue LightSheet (CTLS) system from Intelligent Imaging Innovations was employed for imaging the cleared tissue samples. This system utilizes fiber-coupled lasers with wavelengths of 405, 488, 561, 640, and 785 nm. Beam shaping is handled by a spatial light modulator, and a light sheet is generated using a galvanometric mirror. Illumination objectives (5×/0.14NA) are positioned on both the left and right sides of the imaging chamber, which is filled with BABB solution (refractive index = 1.56). Images are acquired through a Zeiss Plan NeoFluar Z 1×/0.25NA objective lens with a 56 mm working distance. The system integrates a high-speed filter wheel (OPTOSPIN) to rapidly alternate between illumination wavelengths, and an ORCA-fusion BT sCMOS camera is used for detection.

The z-stack was obtained for one wavelength, and the z-stack of the same position was retaken using another wavelength. We chose this strategy because the chromatic focus shift was beyond what the system could dynamically compensate at each z-plane. At each plane, the illumination focus position was shifted three times to virtually create the thin light-sheet. The acquired three images were linearly blended on the fly. Once a z-stack scan was performed, the motorized stage moved the sample in the XY direction to image the next z-stack until the entire tissue volume was covered. Illumination focus was shifted based on the wavelength. Optical zoom of 5× was applied, resulting in pixel sizes of 1.3 μm. A 2 μm z-step was used. All hardware was operated using Slidebook (Intelligent Imaging Innovations), and data was stored on a Dell PowerEdge R740XD with twelve 16 TB hard drives with RAID configuration via a 10-GB Ethernet fiber network.

### Image Analysis

#### Stitching and correcting chromatic aberration

The CTLS produced z-stacks in hdf5 format with xml metafile that can be directly loaded on BigStitcher plugin in Fiji. Using BigStithcer (Hörl et al., 2019), we converted the hdf5 files to n5 with pyramid resolution with the chunk size of 256 × 256 × 128 voxels. The downsampling factor was set to 2 in all dimensions for each pyramid layer and we made up to 5 layers in total. We used BigStitcher for the stitching and the correction of chromatic aberration. In detail, we first performed translational stitching using the channel of rRNA. We used downsampled resolution with the downsampling factor of 16 for the first stitching, and second stitching was done with the downsampling factor of 8. The stitching results were manually curated and corrected until there was no substantial mis-alignment among z-stacks. This step must be extensively and carefully done otherwise will not be able to produce the single-cell precision at the downstream process. We found that the translational stitching with downsampling factor higher than 8 does not help to improve the stitching quality probably because the redundantly high resolution will make the stitching stuck into a local optimal. After the translational stitching, we performed chromatic correction for each z-stack using iterative closest point (ICP) affine transformation with the downsampling factor of 16, 16, and 8 in x, y, and z each and then with the downsampling factor of 8, 8, and 4. The chromatic correction is performed stack by stack, and the results were manually curated and corrected if necessary until there are no substantial mis-alignment. The last step is refinement of the stitching among z-stacks. We used ICP affine stitching across z-stacks while maintaining the relative positions between channels with the downsamling factors of 16, 16 and 8 in x, y and z. If we could not get the single-cell-resolution precision at this step, we cancelled the ICP stitching and went back to the translational stitching until we obtained the single-cell precision. Finally, we performed the ICP affine stitching across z-stacks and channels with the downsampling factors of 8, 8 and 4 in x, y and z. Our visual inspection confirmed this scheme could produce both the stitching and correction of the chromatic aberration at single-cell precision in all the imaging cases. We defined a rectangular boundary to trim off the areas where the tissue does not exist and blended the overlapped regions within the area. We exported 16-bit unsigned integer fused images in n5 format with the chunk size of 256 × 256 × 256 while maintaining the original anisotropic resolution. The resulting n5 contains 5 layers of pyramid resolution with the downsampilng factors of 2 for each layer.

#### Registration of a multi-round imaged sample

We performed 1) global affine registration, 2) global dense deformable registration, and 3) chunk-wise dense deformable registration. The dense deformable registration is based on fast graph-cut optimization with GPU acceleration (Ekström et al., 2020, 2021). The downsampled image (downsampled by 8, 8, and 8 times in x, y, and z) was used for both 1) and 2). The dense intensity-based affine registration was performed with the cross-correlation as a similarity metric. We used the affine transformation matrix for the initial alignment of the global non-linear registration with the cross-correlation as a similarity metric. The obtained displacement field was upsampled for the following chunk-wise registration. Chunk-wise registration is necessary if the data size cannot fit into the system RAM and this makes the strategy appropriate for light-sheet imaging data. We used the higher resolution image (downsampled by 2, 2, and 2 times in x, y, and z) for the chunk-wise non-linear registration. Though we could perform the registration at the original resolution, we did not observe improvements in the quality of the registration. The images were subdivided into chunks with 32 voxels of overlaps. We used the upsampled displacement fields from the previous step as the initial alignment and performed chunk-wise registration. We used cross-correlation as a similarity metric. The obtained chunks of displacement fields were linearly fused and were used to produce the final warped image. The scripts were uploaded to GitHub (https://github.com/tatz-murakami/mFISHwarp).

#### AI-driven segmentation of cells

In the pre-train stage, we adopted a vision transformer (ViT) architecture as a backbone and adopted the masked-autoencoding (MAE) pre-training scheme. One characteristic of images from light-sheet microscopy in cleared tissue is that the discernible features of individual cells are less pronounced, i.e., the shape and texture of each appear remarkably similar, with each resembling a small blob. To better differentiate and segment target cells, we expand the field of view of our *encoder* to include more background context information. Specifically, given a 3D input volume *x* ∈ *R*^*C*×*D*×*H*×*W*^, we define two parts for it: the small center square, referred to as the Region of Interest (ROI), and the input itself, referred to as the context (CON) as showed in **Fig. 4A**. Here, *C* = 2 represents the number of channels, as the input consists of two channels: a reference channel (rRNA) and a signal channel (mRNA). The size of CON, which corresponds to the full input volume *x*, is *C* × *D_*C*ON_*× *H_*C*ON_*× *W_*C*ON_*= 2 × 40 × 1280 × 1280. The ROI is located at the center of the input volume, with a size of *C* × *D_*R*OI_* × *H_*R*OI_*× *W_*R*OI_* = 2 × 40 × 256 × 256. Then, both the ROI and CON are divided into non-overlapped small volumes, named *patches*, of a uniform size *p*^*D*^ × *p*^*H*^ × *p*^*W*^, where 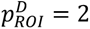,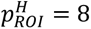,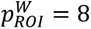 for ROI and 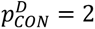,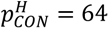,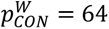 for CON. Pixel values inside a patch are embedded into a *d*-dimensional features using a fully-connected layer followed by adding the *positional embedding*, which is another *d*-dimensional feature depending on the position of the patch inside the whole volume, and a *class embedding*, a *d* -dimensional feature indicating whether the patch belongs to the ROI or CON. This process results in a feature grid of shape 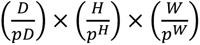, where each element has a dimensionality of *d*. During pre-training, we randomly drop 90% of the patches, flatten the remaining into a 1-dimensional feature sequence, and pass it through a transformer *encoder*, which is later to be used for finetuning. To avoid shortcut learning caused by information leakage, we enforce a mask synchronization rule, i.e., for each patch dropped in CON, patches in ROI that overlaps with the dropped CON patch are also dropped. The encoded features are forwarded through a transformer *decoder* and a pixel reconstruction loss on both the ROI and CON dropped patches. The network is trained in this way for 10 epochs with volumes evenly sampled from two mouse brains with staining of six types of mRNAs.

The second stage is the fine-tuning stage. For simplicity, instead of employing U-Net architectures, we directly concatenate a linear layer after the pre-trained encoder as the segmentation prediction head for fine-tuning the pre-trained model on the cell segmentation task. The pre-trained transformer *encoder* is connected to a randomly-initialized fully-connected layer, projecting features for each patch into the final segmentation prediction of shape 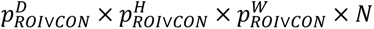. Since the CON is designed to enrich background contextual information, we compute the segmentation loss only within the ROI during fine-tuning. As a result, only patches within the ROI are projected into the final segmentation prediction of shape 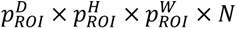, where the *N* is the output channels (**Fig. 4A**). For 2D cell segmentation, *N* = 3, representing the binary probability, x-axis gradients and y-axis gradients, respectively. For 3D cell segmentation, *N* = 4, which includes the binary probability, the x-axis gradients, the y-axis gradients, and the z-axis gradients. More details on the definition and calculation of the x, y, and z gradients can be found in (Stringer et al., 2021). We then aggregate the predictions from all patches by tiling them together to obtain the final output with a shape of *D*_*ROI*_ × *H*_*ROI*_ × *W*_*ROI*_ × *N*.

Given the difficulty of obtaining 3D annotations, we first fine-tune the pre-trained transformer *encoder* using a subset of 2D annotated samples to develop a 2D cell segmentation model. Leveraging the power of pre-training, the 2D cell segmentation model, despite being trained on a limited number of 2D annotated samples, can generate high-quality 3D pseudo-labels that are straightforward for annotators to review and refine. More specifically, given a 3D volume *C* × *D* × *H* × *W*, we first split the volume into *DD* 2D slices, where each slice has a size of *C* × *H* × *W*. Using the fine-tuned 2D cell segmentation model, we generate cell segmentation predictions for each 2D slice. Next, all 2D slices are stitched together to form the 3D pseudo-labels. During this process, cells in one slice are merged with the cells that have the largest overlap area in the adjacent previous slice to ensure continuity across the volume. Subsequently, the pre-trained model is fine-tuned using the refined 3D annotations, resulting in a 3D cell segmentation model capable of generating accurate 3D cell segmentations for entire mouse brains across various genes. During the fine-tuning process, we have 154 annotated 2D slices of size 256 × 256 and 63 annotated 3D volumes of size 80 × 512 × 512 in total. These annotations are randomly selected from 8 genes across three brains: five mouse brains and one human brain.

To demonstrate the advantages of our 3D cell segmentation model, we provide the performance comparison between our model and three other baselines: Cellpose (Stringer et al., 2021), Stardist (Schmidt et al., 2018), Swin UNETR (Hatamizadeh et al., 2022) in **Fig. 4D**. We perform leave-P-out cross-validation using three distinct training-validation dataset splits. Of the 63 annotated 3D volumes, four were generated as pseudo-labels by our fine-tuned 2D cell segmentation model without human proofreading; these were excluded from cross-validation to ensure fairness. To maintain a balanced evaluation, we stratify the training and validation datasets based on the number of cells per image, ensuring a 2:1 ratio in both image count and total cell count. For each training–validation split, all baseline models and our method are fine-tuned on the training dataset and evaluated on the corresponding validation dataset. To achieve the highest-quality whole-brain 3D cell segmentation, we combine the training and validation datasets and retrain our 3D cell segmentation model on the entire dataset.

### Downstream Data Analysis

#### Identification of cells with multiple gene expression

We applied our segmentation model to each channel of the 3D images after registration. To judge whether the segmented object (i.e. cell) expresses multiple genes or not, we calculated the overlapping volume of the segmented object on another object appearing in other channels. If the overlapping volume is more than the threshold, we regarded the objects express multiple genes. We set the threshold of 0.5 for this purpose.

#### Registration to a standard brain atlas

We used Allen Brain Atlas CCFv3 for registration. Using 25-µm voxel-size Nissl-staining template, we registered our rRNA staining 3D image after downsampling. The downsampling was performed so that the voxel size is 32 µm × 20.8 µm × 20.8 µm in z,y, and x each. We used non-linear registration as stated above but without using chunk-wise registration. The resulting displacement field was applied to the centroids of the segmented objects, and the annotation of the region was obtained by referring CCFv3.

## QUANTIFICATION AND STATISTICAL ANALYSIS

### Measurement of phospholipid

The content of the phospholipid in a brain was measured by slightly modifying the method described before (Tainaka et al., 2018). PFA-fixed mouse-brain hemispheres were used for the quantification. The weight of the tissues was measured just after fixation and subjected for the delipidation with MeOH. The delipidated mouse-brain hemispheres were washed with PBS at RT for 1 hour three times. After removing the excess PBS by paper towel, we placed the tissue in a dounce homogenizer. After adding 450 μL of PBS, the tissues were roughly crashed by hand. We added 50 μL of Triton X-100 on top. Using homogenizer 1500 rpm, we completely homogenized the tissue. We collected all the homogenized solution, and PBS was added up to 1 mL. Taking an aliquot of the solution, we measured the phospholipid using Phospholipid Assay Kit (Sigma) by referring to the instruction of the kit. We measured the fluorescence using a plate reader (Cytation1, BioTek).

### Quantification of SNR

To measure the SNR of poly(dT) or NS oligo, we used a hippocampal cell in CA1 as the foreground signal and the non-cellular space of the molecular layer as the background noise. To measure the SNR of HCR amplification using *Gad1*, we used an outer area of the posterior lobe of the cerebellum. The outer area was chosen to avoid the confounding of the penetration of HCR probes. For the signals, we selected Purkinje cells in the areas and measured the averaged intensity. We chose Purkinje cell for this purpose because the cells can be easily identifiable thus removing the confounding of the mixed cell types, and high *Gad1* expression level ensures the quantification of low signals. For the measurement of the noise, we pick up the cells in granular layers except GABAergic Golgi cells and measured averaged intensity. The SNR was obtained by calculating the averaged signals divided by averaged noise.

### Statistical Analysis

Statistical analyses were performed by SciPy. The homogeneous variance for each group was evaluated by Bartlett’s test with a significance level of 0.05. When all of the groups had equal variance, one-way ANOVA, one-way ANOVA with Tukey’s post hoc test, or Student’s t-test was used. When the groups were without equal variance or multiple conditions were grouped for testing, we used the Wilcoxon rank sum test. In this study, p < 0.05 was considered as significant (*p < 0.05, **p < 0.01, ***p < 0.001, and n.s. for not significant evaluations).

## SUPPLEMENTARY NOTE

In organic solvent-based clearing, it is well known that radical generation due to auto-oxidation is a cause of GFP quenching. Dibenzyl ether, as demonstrated by Becker et al. (Becker et al., 2012), produces peroxides through auto-oxidation; thus, alumina purification is performed prior to its use. Among the components of BABB, benzyl benzoate has been employed as a clearing solvent without the need for pre-purification to remove peroxides (Jing et al., 2018), suggesting that auto-oxidation is effectively suppressed. However, benzyl alcohol decomposes slowly to benzaldehyde when exposed to air (Kulkarni and Mehendale, 2005), which is accompanied by radical formation in the system. Therefore, alternative compounds with resistance to auto-oxidation were investigated as substitutes for benzyl alcohol. Although benzyl benzoate—like benzyl alcohol and dibenzyl ether—contains a benzyl carbon that serves as a potential radical generation site, the oxygen atom adjacent to the benzyl carbon is part of an ester group, and this structural feature is expected to suppress radical formation. Consequently, benzyl acetate, which also contains an ester moiety and is miscible with hydrogen peroxide, was selected for further evaluation.

To theoretically assess whether these compounds exhibit oxidation resistance comparable to that of benzyl benzoate, quantum chemical predictions were performed for benzyl alcohol, benzyl benzoate, dibenzyl ether, and benzyl acetate. First, Fukui indices (specifically, f⁻) were calculated as a conceptual reactivity analysis to identify the sites most prone to initiating radical formation. The results indicated that, among the four chemicals, the most electrophilically reactive sites— potential initiation points for radical generation—were located on the benzyl carbons.

Next, because the actual formation of radicals requires homolytic cleavage of the C–H bond at the benzyl carbon, the bond dissociation energies (BDE) of the corresponding C–H bonds were evaluated. The computed BDE values were as follows: benzyl alcohol = 3.678 eV, benzyl benzoate = 3.821 eV, dibenzyl ether = 3.676 eV, and benzyl acetate = 3.804 eV, indicating that benzyl acetate exhibits a BDE comparable to that of benzyl benzoate.

Furthermore, a Natural Bond Orbital (NBO) analysis was carried out to quantify the extent of electron donation from the lone pairs of the oxygen atoms to the σ* orbital of the benzyl C–H bond—a phenomenon attributable to hyperconjugation or dipole interactions—by evaluating the second-order perturbation energies, E(2) (in kcal/mol). The values obtained were: benzyl alcohol = 6.69, benzyl benzoate = 5.53, dibenzyl ether = 7.14, and benzyl acetate = 5.50. The higher E(2) values in benzyl alcohol and dibenzyl ether reflect increased electron donation from oxygen, thereby facilitating radical generation. In contrast, the E(2) value of benzyl acetate, which is similar to that of benzyl benzoate, suggests that electron donation from oxygen is suppressed and that there are fewer factors promoting radical formation.

Taken together, these results support the evaluation of benzyl acetate as an oxidation-resistant solvent, comparable to benzyl benzoate, for use in organic solvent-based tissue clearing applications.

## Methods

All quantum chemical calculations were carried out using ORCA (Program Version 5.0.4) at the B3LYP/def2 - TZVP level of theory. The ground-state geometries of the studied molecules were fully optimized prior to subsequent analyses. To evaluate the tendency for radical formation, Fukui indices were computed using a finite difference method. In particular, the electrophilic Fukui index (f⁻) was determined from the difference in electron populations at the reactive benzyl sites between the neutral and the corresponding cation species, i.e., f⁻ = q(neutral) – q(cation) (Fukui, 1982; Yang and Mortier, 1986). (Note: This finite difference approximation at the atomic level quantifies the propensity of each atom to undergo oxidation, a descriptor widely used to predict local reactivity.) Additionally, bond dissociation energies (BDEs) for the benzylic C–H bonds were calculated using the equation: BDE = E(radical) + E(H) − E(parent), where E(radical) is the energy of the radical species formed upon homolytic cleavage, E(H) is the energy of the isolated hydrogen atom, and E(parent) is the energy of the optimized parent molecule. Zero-point energy corrections obtained from frequency calculations were applied to all species. Natural Bond Orbital (NBO) analysis was also performed to quantify the second-order perturbation energies associated with electron donation from adjacent oxygen lone pairs to the antibonding C–H σ* orbitals (Reed et al., 1985; Neese, 2012).

## DATA AND SOFTWARE AVAILABILITY

Our Python codes for ZenCell is available at https://github.com/mx41-m/ZenCell.

## RESOURCE TABLE

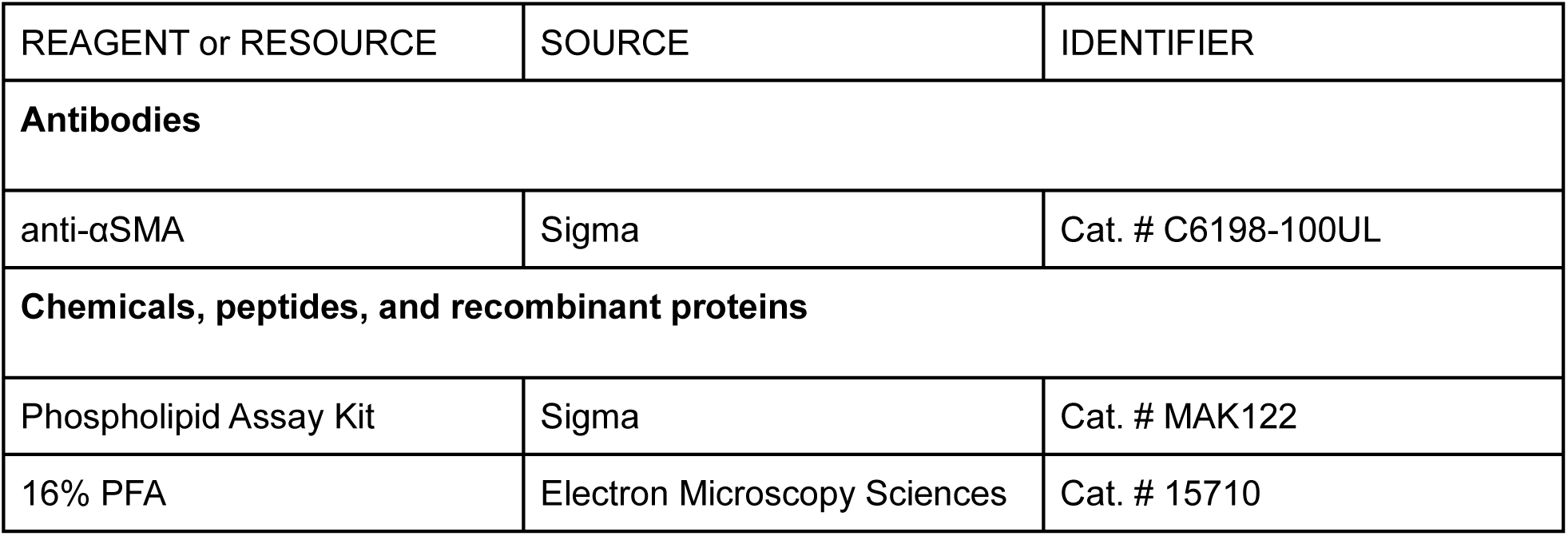
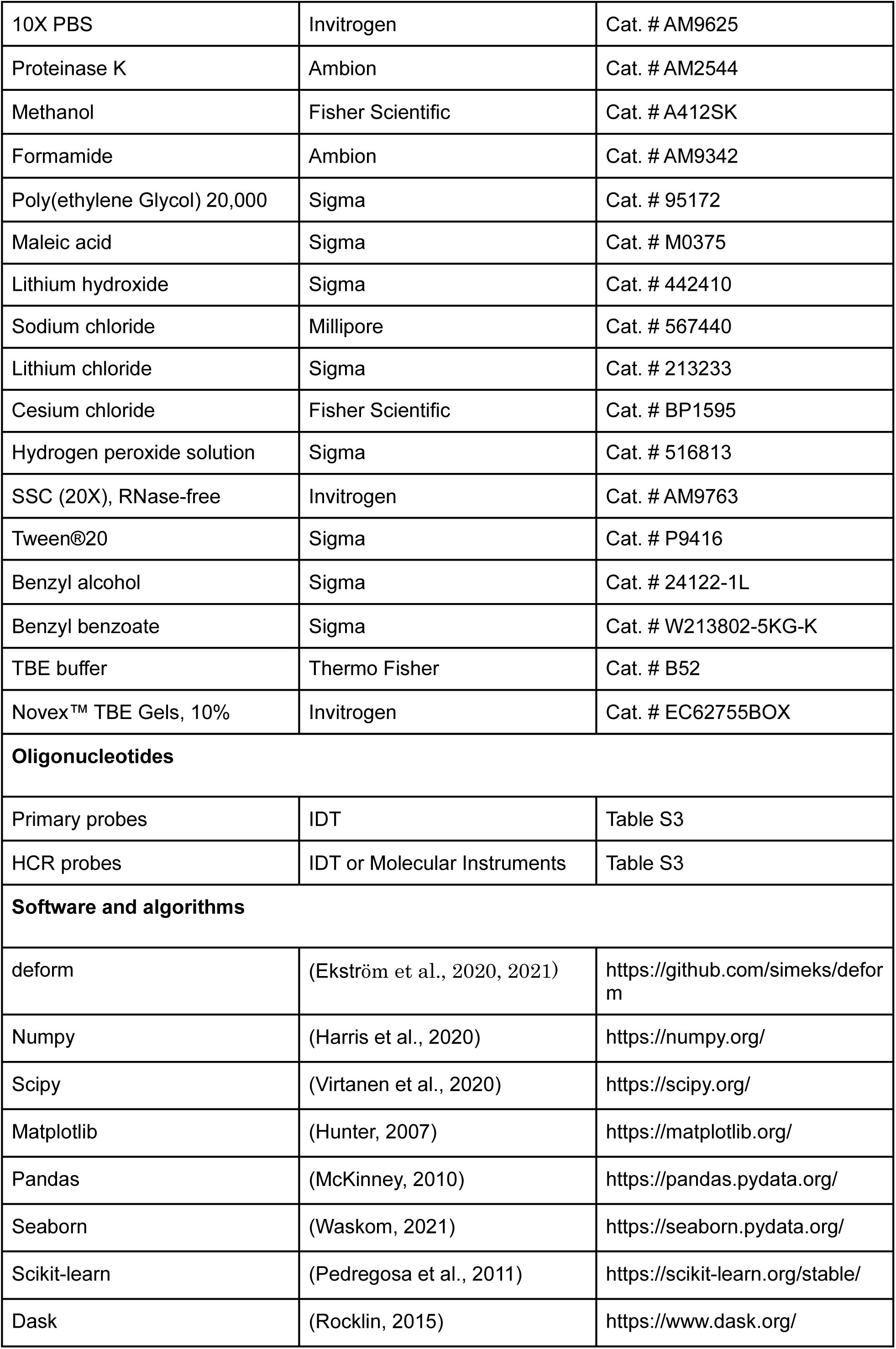
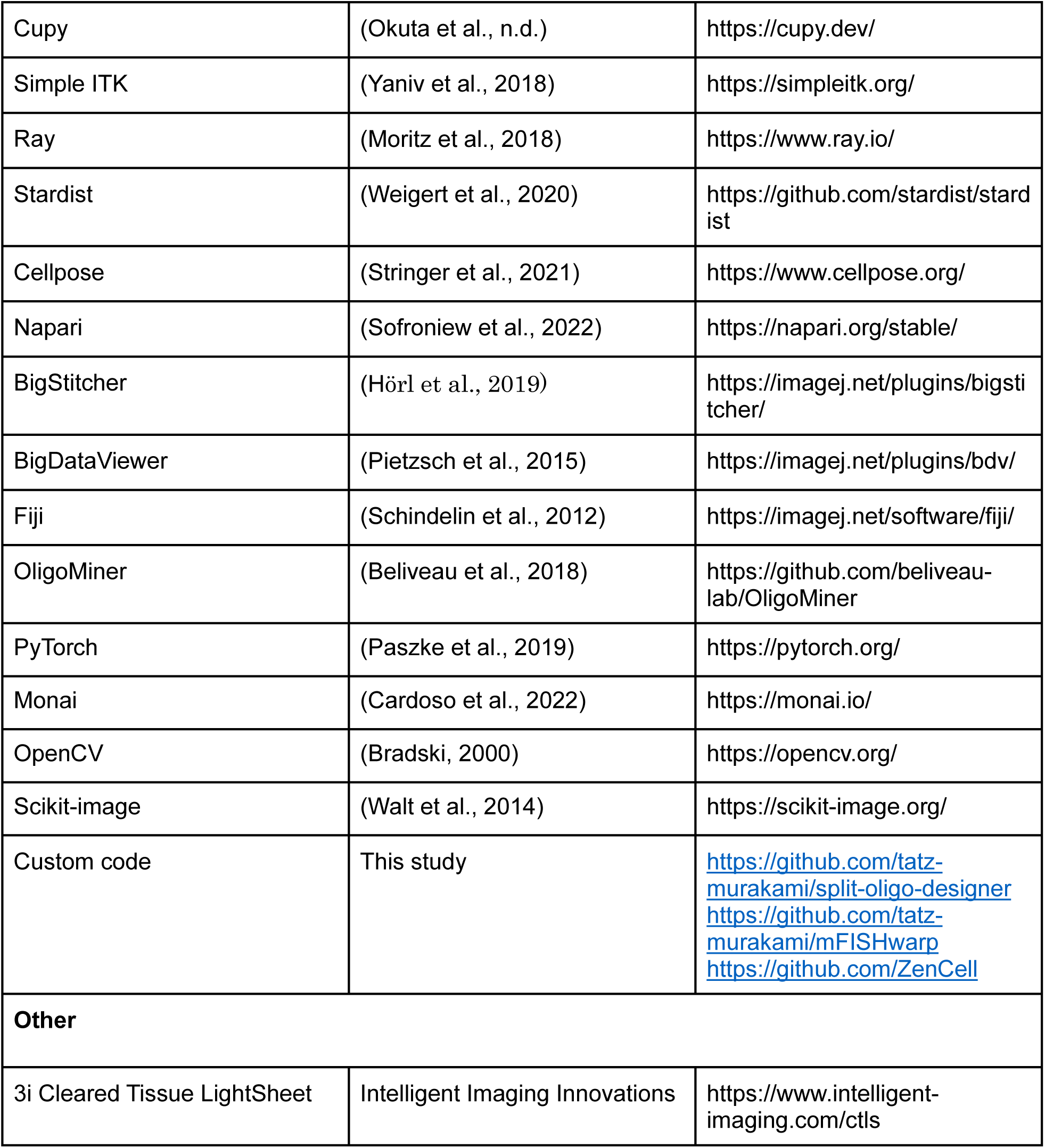

| REAGENT or RESOURCE | SOURCE | IDENTIFIER |
| --- | --- | --- |
| **Antibodies** |  |  |
| anti-αSMA | Sigma | Cat. # C6198-100UL |
| **Chemicals, peptides, and recombinant proteins** |  |  |
| Phospholipid Assay Kit | Sigma | Cat. # MAK122 |
| 16% PFA | Electron Microscopy Sciences | Cat. # 15710 |
| 10X PBS | Invitrogen | Cat. # AM9625 |
| Proteinase K | Ambion | Cat. # AM2544 |
| Methanol | Fisher Scientific | Cat. # A412SK |
| Formamide | Ambion | Cat. # AM9342 |
| Poly(ethylene Glycol) 20,000 | Sigma | Cat. # 95172 |
| Maleic acid | Sigma | Cat. # M0375 |
| Lithium hydroxide | Sigma | Cat. # 442410 |
| Sodium chloride | Millipore | Cat. # 567440 |
| Lithium chloride | Sigma | Cat. # 213233 |
| Cesium chloride | Fisher Scientific | Cat. # BP1595 |
| Hydrogen peroxide solution | Sigma | Cat. # 516813 |
| SSC (20X), RNase-free | Invitrogen | Cat. # AM9763 |
| Tween®20 | Sigma | Cat. # P9416 |
| Benzyl alcohol | Sigma | Cat. # 24122-1L |
| Benzyl benzoate | Sigma | Cat. # W213802-5KG-K |
| TBE buffer | Thermo Fisher | Cat. # B52 |
| Novex™ TBE Gels, 10% | Invitrogen | Cat. # EC62755BOX |
| **Oligonucleotides** |  |  |
| Primary probes | IDT | Table S3 |
| HCR probes | IDT or Molecular Instruments | Table S3 |
| **Software and algorithms** |  |  |
| deform | (Ekström et al., 2020, 2021) | https://github.com/simeks/deform |
| Numpy | (Harris et al., 2020) | https://numpy.org/ |
| Scipy | (Virtanen et al., 2020) | https://scipy.org/ |
| Matplotlib | (Hunter, 2007) | https://matplotlib.org/ |
| Pandas | (McKinney, 2010) | https://pandas.pydata.org/ |
| Seaborn | (Waskom, 2021) | https://seaborn.pydata.org/ |
| Scikit-learn | (Pedregosa et al., 2011) | https://scikit-learn.org/stable/ |
| Dask | (Rocklin, 2015) | https://www.dask.org/ |
| Cupy | (Okuta et al., n.d.) | https://cupy.dev/ |
| Simple ITK | (Yaniv et al., 2018) | https://simpleitk.org/ |
| Ray | (Moritz et al., 2018) | https://www.ray.io/ |
| Stardist | (Weigert et al., 2020) | https://github.com/stardist/stardist |
| Cellpose | (Stringer et al., 2021) | https://www.cellpose.org/ |
| Napari | (Sofroniew et al., 2022) | https://napari.org/stable/ |
| BigStitcher | (Hörl et al., 2019) | https://imagej.net/plugins/bigstitcher/ |
| BigDataViewer | (Pietzsch et al., 2015) | https://imagej.net/plugins/bdv/ |
| Fiji | (Schindelin et al., 2012) | https://imagej.net/software/fiji/ |
| OligoMiner | (Beliveau et al., 2018) | https://github.com/beliveau-lab/OligoMiner |
| PyTorch | (Paszke et al., 2019) | https://pytorch.org/ |
| Monai | (Cardoso et al., 2022) | https://monai.io/ |
| OpenCV | (Bradski, 2000) | https://opencv.org/ |
| Scikit-image | (Walt et al., 2014) | https://scikit-image.org/ |
| Custom code | This study | https://github.com/tatz-murakami/split-oligo-designer<br>https://github.com/tatz-murakami/mFISHwarp<br>https://github.com/ZenCell |
| **Other** |  |  |
| 3i Cleared Tissue LightSheet | Intelligent Imaging Innovations | https://www.intelligent-imaging.com/ctls |

## ACKNOWLEDGEMENTS

This work was supported by the Howard Hughes Medical Institute (to N.H.), CHDI Foundation (to N.H.), Fisher Center for Alzheimer’s Research Foundation (to N.H.), Fisher AI Platform (to N.H.), Japanese Society for Promotion of Science Overseas Research Fellowship (to T.C.M), and Leon Levy Scholarships in Neuroscience (to T.C.M.). We thank Christina Pressl, Katherine Kane, Laura Kus, and Maytal Babajanian for their technical assistance in the experiment on human tissues. The human tissues presented in this study were provided by Miami’s Endowment Brain Bank. The mice were bred in Hatten lab at Rockefeller University and provided by Yin Fang and Zack Schapire, and mice were perfused by Xing Jie. Yuchen Zhang helped generating training dataset for cell segmentation. We thank Balakanagaram Jayaraman, Jason Banfelder, Nicholas Didkovsky, and Rebecca Bennett for their support in setting up the infrastructure of the data storage. We thank Etsuo A. Susaki for providing with *Fos* immunostaining dataset.

## AUTHOR CONTRIBUTIONS

T.C.M. designed the study. T.C.M. developed mFISH3D. T.C.M. and Y.M performed the experiments. M.X and Z.L. developed ZenCell and M.X. implemented it. Y.Y. developed the manual annotation tool and examined the performance of ZenCell.

T.M. and S.R collected the squid brains and designed oligonucleotides for squid tissue and analyzed the images of a squid brain. K.T. proposed the safer alternative to benzyl alcohol and conducted quantum chemistry analysis. T.C.M. performed the image analysis. T.C.M and N.H. supervised the research. M.X supervised deep learning. T.C.M drafted the manuscript, and all authors reviewed and edited the manuscript. All authors discussed the results and commented on the manuscript.

N.H. acquired research funding.

## LEAD CONTACT

Further information and requests for resources and reagents should be directed to and will be fulfilled by the Lead Contact.

## SUPPLEMENTARY FIGURE TITLES AND LEGENDS

**Figure S1.**
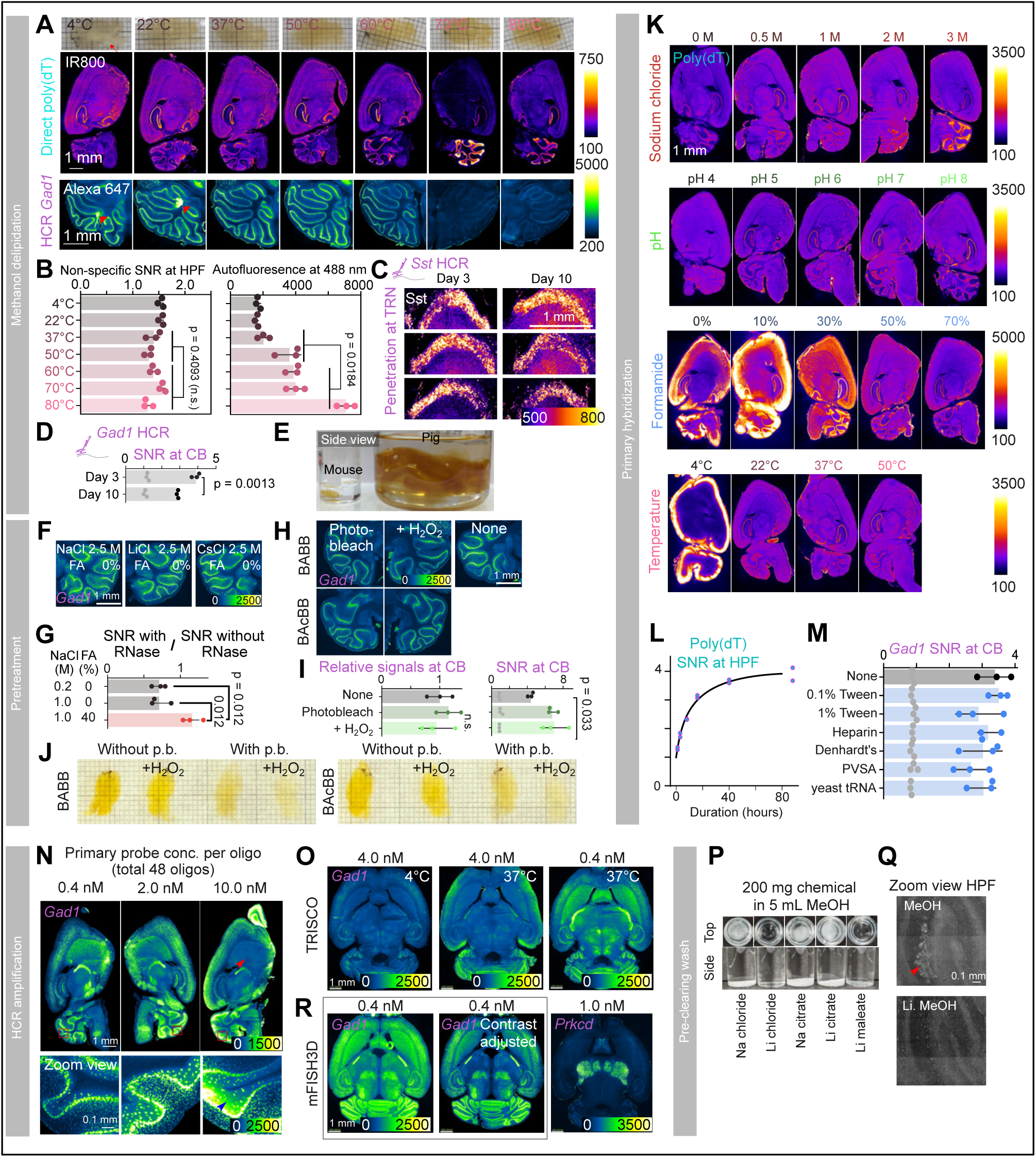
Bottom-up design of 3D fluorescent in situ hybridization. (**A**) Clearing performance and FISH signals after MeOH delipidation at different temperatures. (Top) Photographs of brains after clearing. (Middle) Representative sagittal z-plane of the staining of poly(dT). (Bottom) Representative sagittal z-plane of the staining of *Gad1* HCR in cerebellums. (**B**) (Left) Quantification of SNR of non-specific binding at HPF. Data are mean ± SD (N = 3). Wilcoxon rank sum test was used. (Right) Quantification of autofluorescence at HPF. Data are mean ± SD (N = 3). Wilcoxon rank sum test was used. (**C**) Penetration of HCR probes against *Sst* after delipidation with 3 or 10 days. (**D**) Quantification of SNR of *Gad1* HCR after delipidation with 3 or 10 days. Data are mean ± SD (N = 3). Student’s t-test was used. (**E**) Side-view photography of cleared pig brain hemisphere and whole-mouse brain. (**F**) Gad1 HCR staining after pre-treatment with different types of salts. (**G**) Quantification of SNR after pre-treatment with RNase in various compositions of buffer. Data are mean ± SD (N = 3). One-way ANOVA with Tukey’s post hoc test was used. (**H**) Gad1 HCR staining after photobleaching or H2O2-assisted photobleaching. BAcBB: a mixture of benzyl acetate and benzyl benzoate. Images were taken in BABB for comparison. See also **Methods**. (**I**) (Left) Relative signal intensity of *Gad1* HCR in the cerebellum after photobleaching. The mean value of signals without photobleaching was set to one. (Right) Quantification of SNR of *Gad1* HCR in cerebellum. Data are mean ± SD (N = 3). One-way ANOVA with Tukey’s post hoc test was used. (**J**) Bleaching performance of hydrogen peroxide BAcBB. Photos were taken after replacing the solvent with BABB for comparison. (**K**) Representative z-planes of poly(dT) staining in various chemical conditions. (**L**) Time-course quantification of SNR of poly(dT) (N = 2). Polynomial curve fitting is shown. (**M**) Quantification of SNR of *Gad1* HCR after primary hybridization with various additive chemicals. 50 µg/ml of heparin, 1× concentration of Denhardt’s solution, 0.3% of poly(vinylsulfonic acid) (PVSA), and 0.5 mg/ml yeast tRNA were used. Data are mean ± SD (N = 3). (**N**) Representative z-planes of *Gad1* HCR after primary hybridization with various concentrations of primary probes. (Top) Overviews. The red arrowhead indicates the area where the penetration of HCR probes is limited. (Bottom) Magnified views. The blue arrowhead indicates the area with non-specific amplification. (**O**) TRISCO with different HCR reaction temperatures and varying concentrations of primary probes. The molar values indicate the concentration of primary probes per oligonucleotide. (**P**) Solubility test of the chemicals in methanol. Top views and side views are shown. Na: sodium. Li: lithium. (**Q**) Representative z-planes in HPF after pre-clearing wash with lithium chloride in methanol. The red arrowhead indicates the deposits. (**R**) Whole-mouse-brain in situ hybridization against *Gad1* and *Prkcd*. Imaging was performed under conditions designed to allow comparison with **O**. The molar values indicate the concentration of primary probes per oligonucleotide. For **A**, **C**, **F**, **H**, **K**, **N**, **O**, and **R**, the intensity of the images is color-coded according to the bars.

**Figure S2.**
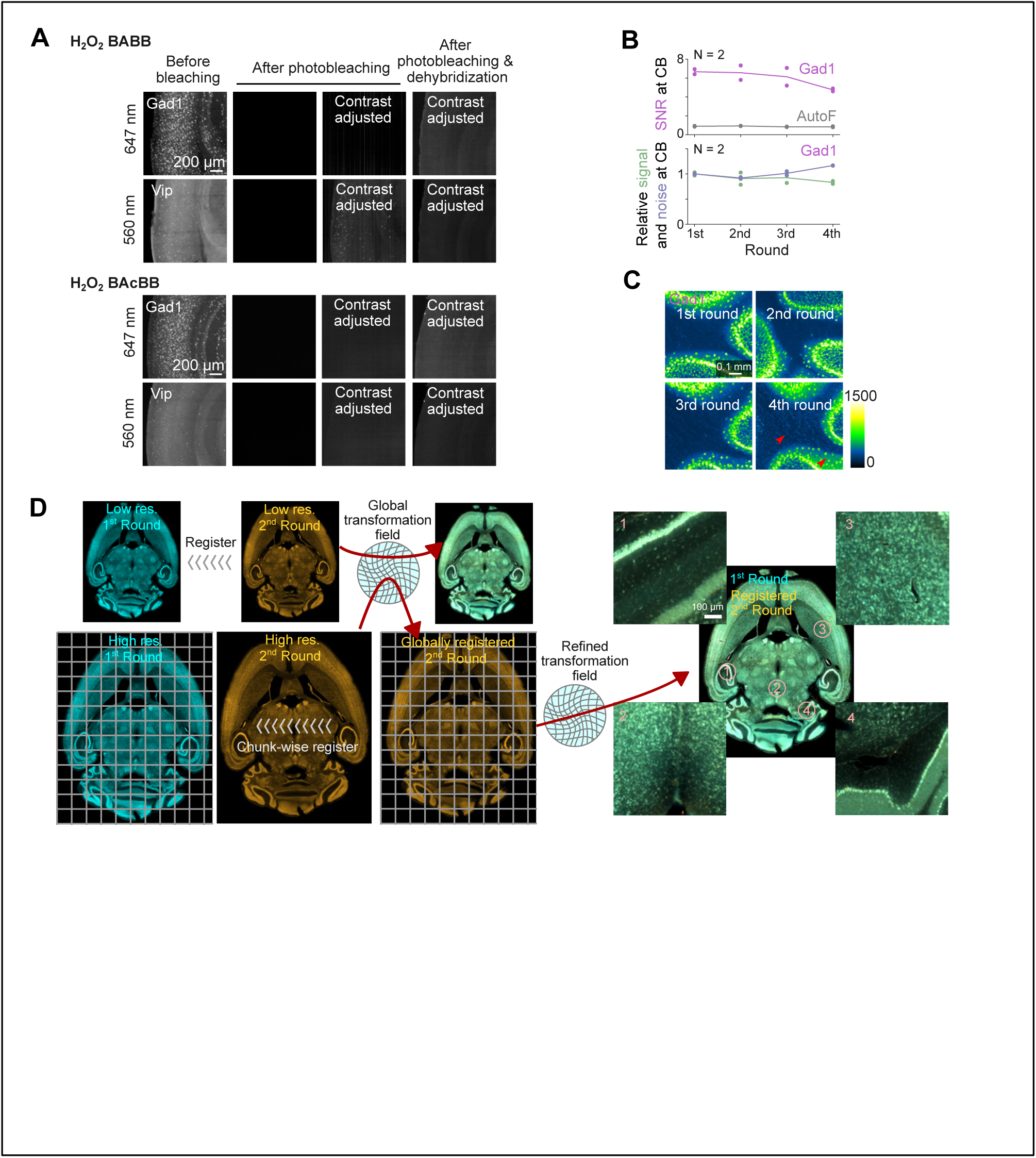
Multiplexed fluorescent in situ hybridization (mFISH3D) in a whole-mouse brain. (**A**) The effect of H2O2-assisted photobleaching and dehybridization to quenches the fluorescent labels. Magnified views of representative z-planes are shown. Top: photobleaching with hydrogen peroxide BABB. Bottom: photobleaching with hydrogen peroxide BAcBB. (**B**) (Top) Quantification of SNR of *Gad1* HCR at cerebellum after multiple rounds of H2O2-assisted photobleaching and dehybridization. SNR of autofluorescence is also included as a control. (Bottom) Signal and noise of *Gad1* HCR. (**C**) Magnified representative z-planes at cerebellum of Gad1 HCR after multiple rounds of H2O2-assisted photobleaching and dehybridization. (**D**) Schematic illustration of single-cell-resolution image registration.

**Figure S3.**
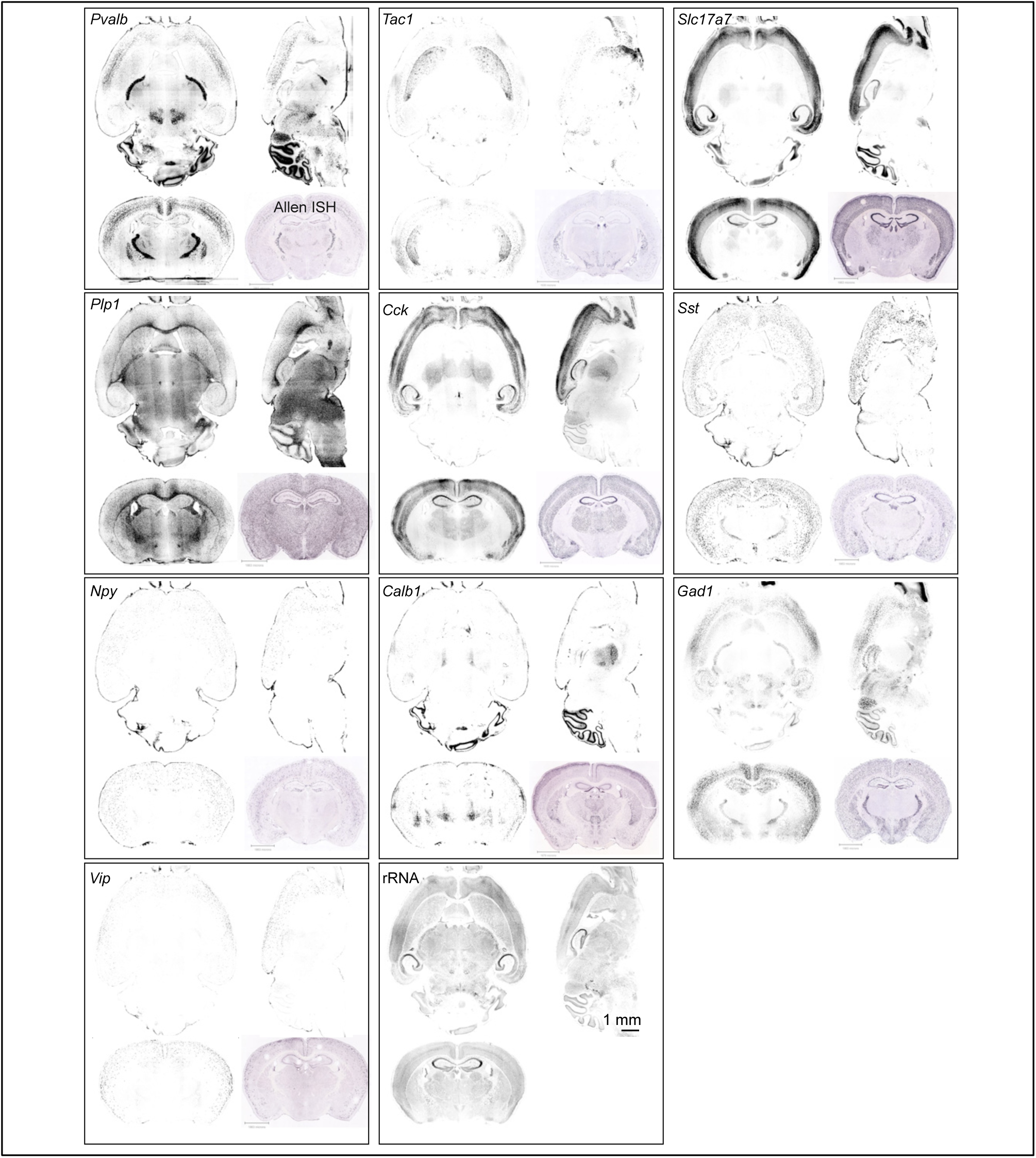
Multiplexed fluorescent in situ hybridization (mFISH3D) in a whole-mouse brain. Orthogonal views. Horizontal, sagittal and coronal views are shown along with in situ hybridization images from Allen Brain Institute (Allen ISH).

**Figure S4.**
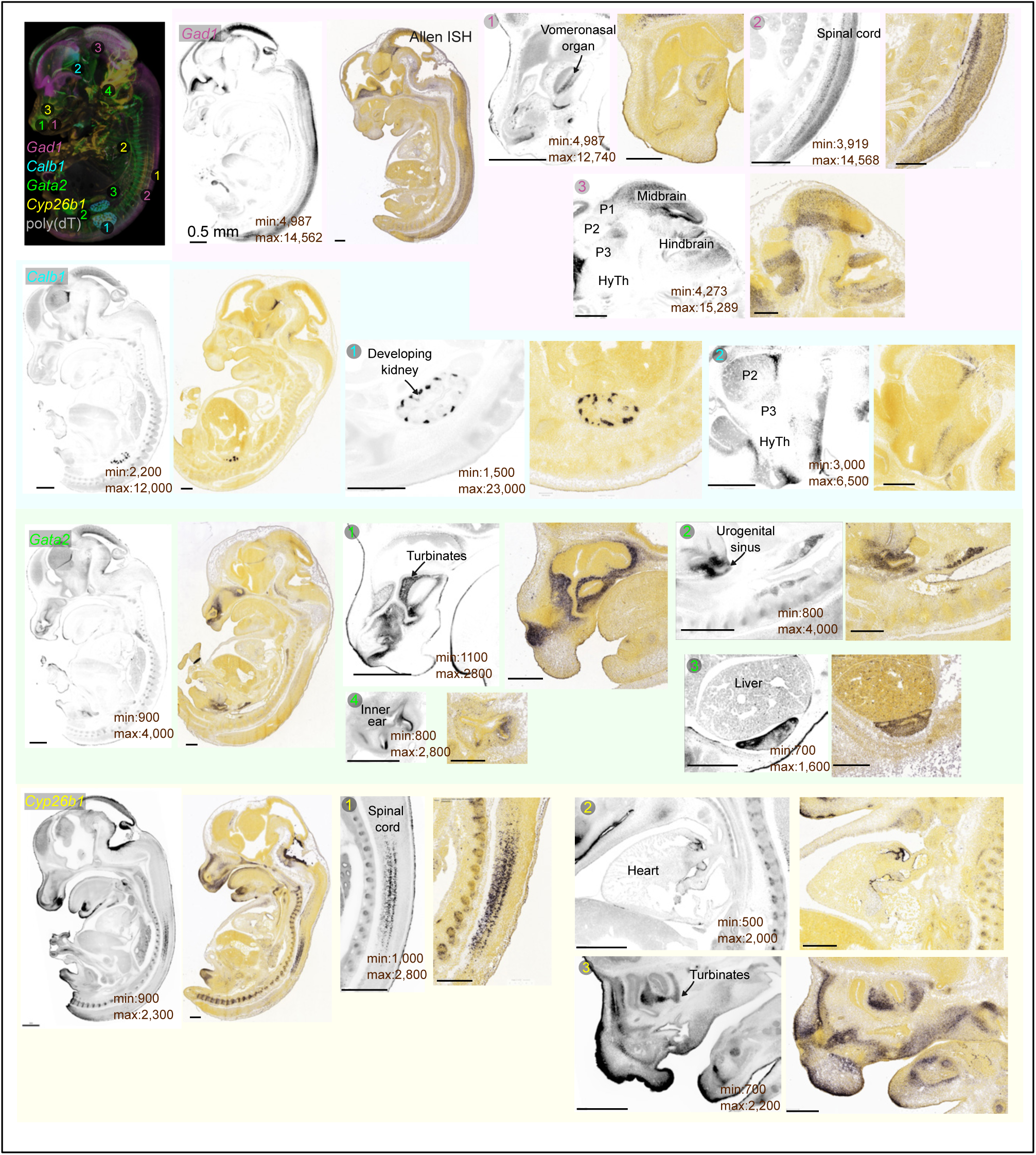
Application of mFISH3D in E13.5 mouse embryo. Representative z-planes are shown to demonstrate the specificity of the staining. The contrast of the images was adjusted for each for the visualization purpose. Volumetric rendering is shown in the top left corner to indicate the positions shown in each panel. P1-3: prosomere 1-3, HyTh: hypothalamus. Scale bars are 0.5 mm. For reference, we included the chromogenic ISH images from Allen Brain Institute.

**Figure S5.**
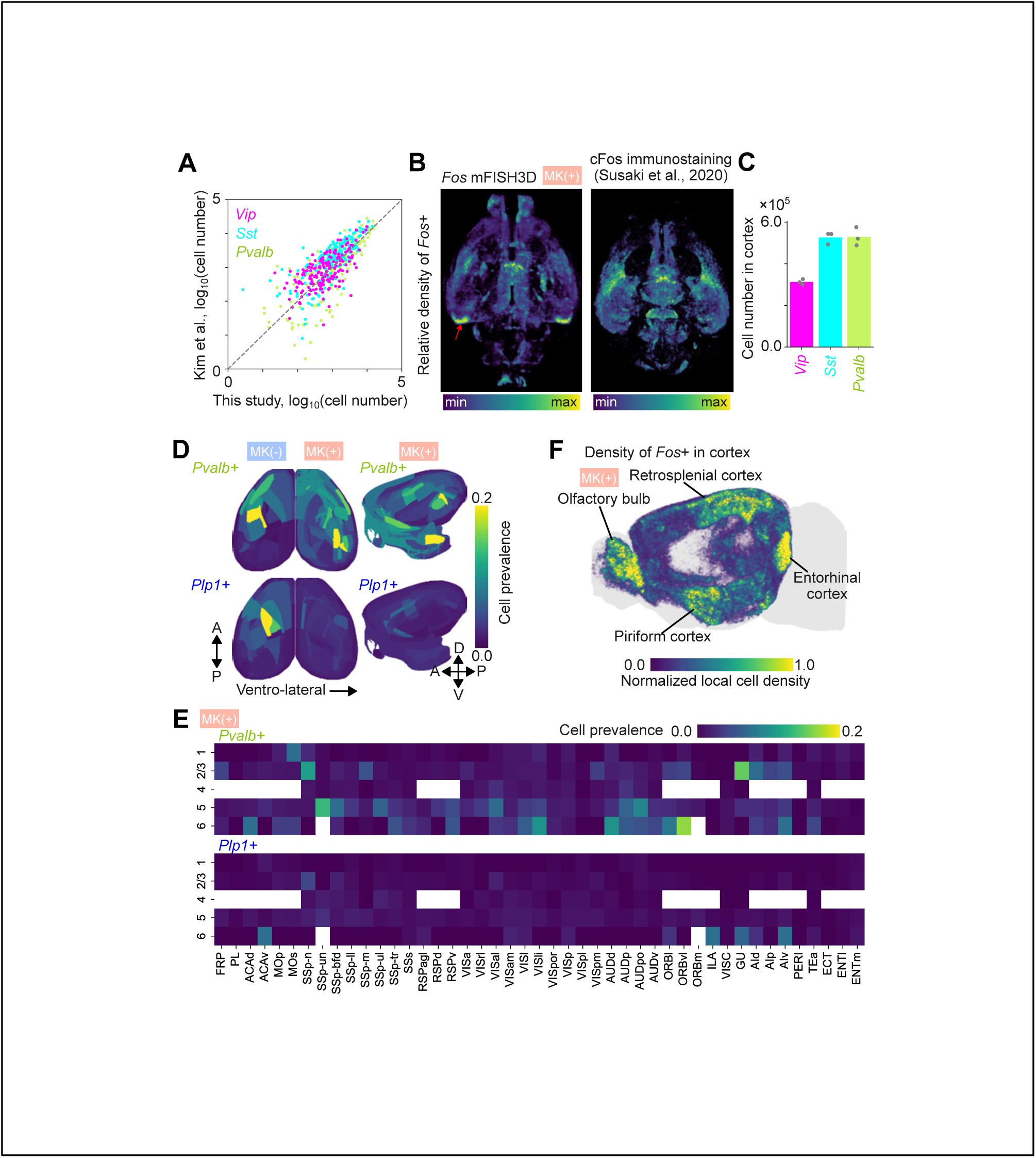
Artificial intelligence-driven segmentation of mFISH3D using ZenCell. (**A**) Correlation plot of the number of cortical inhibitory neurons between this study and the study by Kim et al., 2017. Each dot represents an anatomical region as specified in Allen Brain Atlas CCFv3. (**B**) Relative density of *Fos*+ cells. The densities were locally calculated and globally normalized by min-max normalization. The red arrow indicates the entorhinal cortex, where mRNA expression is more explicit than protein. (**C**) Total cell number by cell types in cortex. (N=3) (**D**) Spatial mapping of cell prevalence in *Fos*+ cells of *Pvalb*+ or *Plp1*+. Left: horizontal projection. Right: sagittal projection. (**E**) Heatmap of cell prevalence of *Pvalb*+ or *Plp1*+ in isocortex by layers. (**F**) Density mapping of *Fos+* cells in cortex. Each dot indicates the position of the cells. Local cell density was calculated and shown in color.

**Figure S6.**
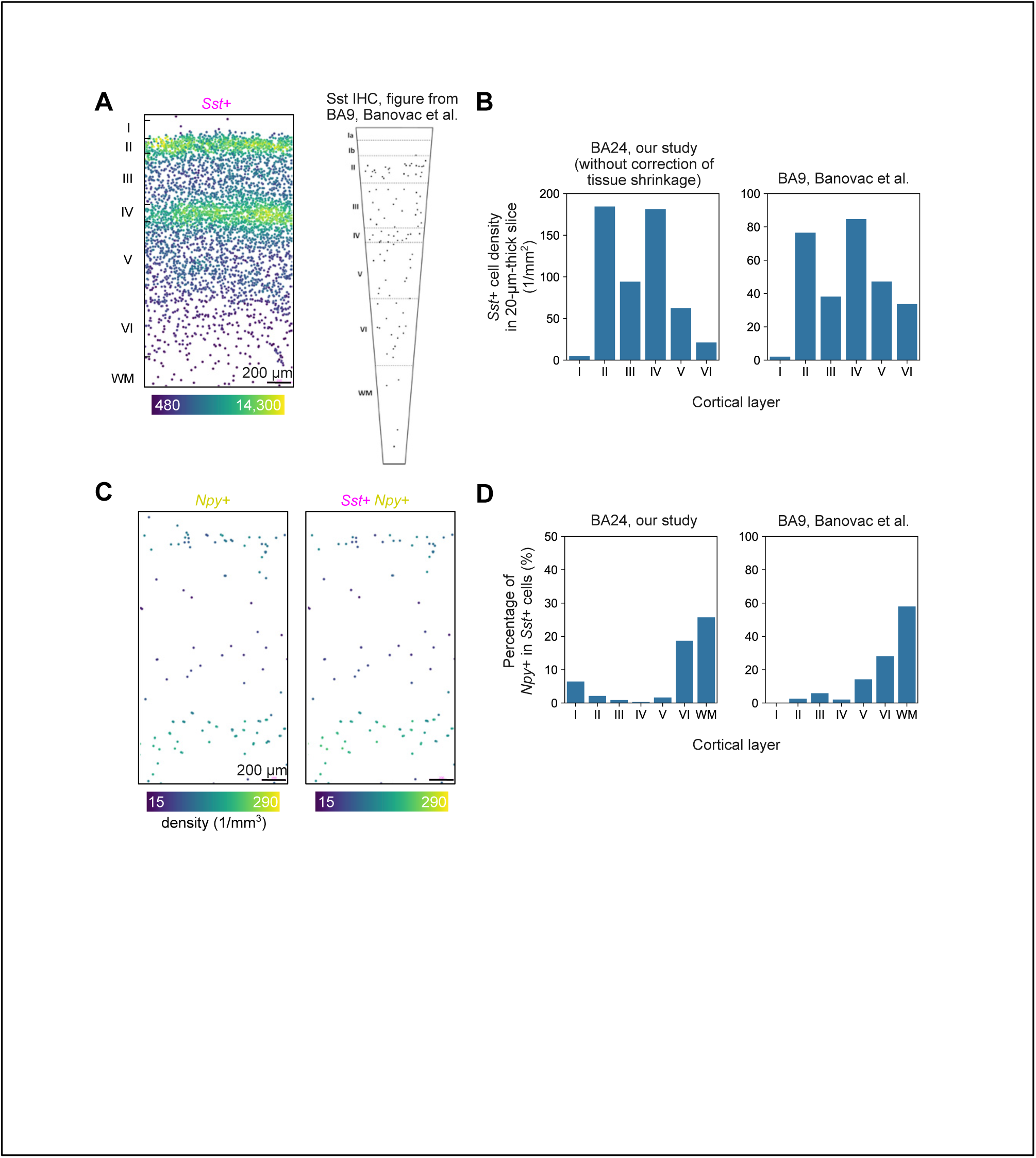
Laminar distribution of *Sst*+ and *Npy*+ cells in human cortex. (**A**) Left: Positions of *Sst*+ cells in the cortex. Each dot represents the position, and color represents the local cell density. Z-projection of a 420-µm-thick volume is shown for visualization purpose. Right: Position of *Sst*+ cells in BA9 cortex using immunohistochemistry. Figure from Banovac et al. (**B**) *Sst*+ cell density by layer. The density was recalculated to indicate the area density in a 20-µm-thick slice. The volume change caused by tissue shrinkage during tissue clearing was not used to correct the density. (**C**) Left: Positions of *Npy*+ cells. Right: Positions of *Sst*+*Npy*+ cells. Each dot represents the position, and color represents the local density. Z-projection of a 420-µm-thick volume is shown for visualization purpose. (**D**) Percentage of *Npy*+ cells in *Sst*+ cells by layer. For **C** and **D**, the data shown on the right were obtained from Banovac et al.

## SUPPLEMENTARY TABLE AND VIDEO INFORMATION

**Supplementary Video 1.** Serial lateral images of mouse brains after staining against *Gad1* or *Prkcd*.

**Supplementary Video 2**. Serial lateral images of an rRNA-stained mouse brain after single-cell-resolution image registration.

**Supplementary Video 3**. Serial lateral images of a mouse brain after mFISH3D against 10 types of mRNA, rRNA and IHC against αSMA.

**Supplementary Video 4**. Serial lateral images of a mouse brain after mFISH3D against 10 types of mRNA with magnified views. Insets show magnified views with locally adjusted contrast for the better visibility.

**Supplementary Video 5**. Volume rendering and serial lateral images of a mouse embryo after mFISH3D. The signal at the tissue rim was masked in the volume rendering for better visualization.

**Supplementary Video 6**. Volume rendering and serial lateral images of an adult mouse kidney after mFISH3D.

**Supplementary Video 7**. Segmentation of a whole brain with ZenCell. The original image, the segmentation overlaid on the image, and the segmentation are shown.

**Supplementary Video 8**. Volume rendering and serial lateral images of a piece of human cingulate cortex.

